# From co-occurrence to increasingly precise temporal coordination: Development of slow oscillation–spindle coupling from infancy to toddlerhood

**DOI:** 10.64898/2026.08.30.748102

**Authors:** Eva-Maria Kurz, Matthias Mölle, Jan Born, Manuela Friedrich

**Affiliations:** Institute of Medical Psychology and Behavioral Neurobiology, University of Tübingen, Tübingen, Germany; Department of Child and Adolescent Psychiatry, Psychosomatics and Psychotherapy, University Hospital of Psychiatry and Psychotherapy, University of Tübingen, Tübingen, Germany; Department of Psychology, Development and Neurodiversity Lab, Uppsala University, Uppsala, Sweden; Center of Brain, Behavior and Metabolism (CBBM), University of Lübeck, Lübeck, Germany; Institute for Diabetes Research and Metabolic Diseases of the Helmholtz Center Munich, University Tübingen (IDM), Tübingen, Germany; Center for Integrative Neuroscience, University of Tübingen, Tübingen, Germany; German Center for Diabetes Research (DZD), Tübingen, Germany; German Center for Mental Health (DZPG), Tübingen, Germany; Department of Psychology, Humboldt-University Berlin, Berlin, Germany

**Keywords:** Development, slow oscillation, sleep spindles, slow oscillation-spindle coupling, infants, toddlers

## Abstract

The precise temporal coordination of sleep spindles and slow oscillations (SOs) is a key mechanism of memory consolidation during sleep, yet little is known about its emergence early in life. Here, we investigated the development of SO-spindle coupling in 9-to 16-monthold infants and toddlers. While morphologies of spindles and SOs were largely comparable between infants younger than one year and toddlers older than one year, their temporal coordination showed clear age-related changes. In infants, SO-spindle co-occurrence was highest over occipital cortex regions, as was to be expected by chance based on the spatial distribution of the two oscillations. Co-occurrence rates in infants already exceeded chance level, but only at central regions where the highest co-occurrence rates were observed in toddlers. Moreover, a nesting of spindles into the upstate of SOs was evident in fronto-central regions even in infants, and increased in strength and precision with age. Together, these findings suggest that coordinated SO-spindle coupling emerges in the second half of the first year of life and becomes increasingly precise across the transition into toddlerhood.

## Introduction

Sleep changes rapidly in the first two years of life. This maturation is observed on a macro scale reflected by the transition from several shorter sleep bouts across the day to longer sleep periods during the night (e.g. Galland et al., 2012) but also on a micro-scale reflected by the developmental changes of sleep oscillations. The hallmarks of non-rapid eye movement (non-REM) sleep, slow oscillations (SOs) and sleep spindles follow distinct developmental trajectories. SOs (~ 1 Hz), which originate in the neocortex and reflect alternated hyper- and depolarization of underlying neuronal membrane potentials (Nir et al., 2011; Steriade et al., 2001), show highest density at posterior regions during early life (Page et al., 2018) with a shift to the frontal cortex across childhood (Kurth et al., 2010). Spindle rates are highest at frontal and central areas during infancy, showing a shift to central regions during toddlerhood (Kwon et al., 2023).

While spindles have repeatedly been associated with memory consolidation in infants, toddlers, and older children (Bastian et al., 2024; Friedrich et al., 2022; Friedrich et al., 2019, 2020; Friedrich et al., 2015; Hahn et al., 2019; Kurdziel, 2019), it is their coordinated co-occurrence together with SOs and hippocampal ripples that is thought to promote active systems consolidation in the mature brain (Klinzing et al., 2019). Ripples that are accompanied by the replay of neuronal cell assemblies are nested within spindle throughs which themselves are nested within the excitable upstate of SOs, thus providing a milieu for the transmission and integration of hippocampal information into neocortical longterm stores (Brodt et al., 2023; Staresina et al., 2015). The precise coupling of spindles towards the SO upstate benefits memory consolidation in both adults and school-aged children (Ng et al., 2025). Across childhood, the coupling of spindles to SOs has shown to increase in precision (Hahn et al., 2020; Kurz et al., 2023), with this mechanism possibly being dependent on the maturation of sleep spindles to adult-like patterns (Joechner et al., 2023).

To the best of our knowledge, so far, only two studies explored the temporal coordination of spindles with SOs in infants. In 6-months-olds, greatest co-occurrence rates have been observed in occipital areas (Jaramillo et al., 2023). In our own study (Kurz et al., 2024), co-occurrence rates did not exceed chance level in two to three months old infants at any site, and there was also no evidence to suggest that sleep spindles and SOs occur temporally coordinated at this age. In contrast, both above-chance co-occurrence of spindles and SOs and the coupling of spindles to the upstate of SOs were present at frontal and central regions in 14 to 17 months old toddlers (Kurz et al., 2024). In this study, we did not record occipital sites.

There are currently significant gaps in our knowledge regarding the development of the temporally coordinated occurrence of sleep spindles and SOs in infants and the emergence of the precise nesting of spindles into the upstate of SOs. To fill these gaps, we investigated the coordinated coalescence of spindles and SOs in a sample of 9 to 16 months old infants. We expected older participants to exhibit greater co-occurrence rates than younger ones and hypothesized that infants younger than one year already show above chance level co-occurrence of spindles and SOs. Having now recorded brain activity also at occipital sites, we specifically aimed to investigate whether SO-spindle co-occurrence in the younger participants exceeds chance level at occipital regions. Finally, we hypothesized that the temporally precise coupling of sleep spindles with SOs emerges within the investigated age range and increases with increasing age.

## Methods

### Participants

We analyzed sleep EEG recordings from a nap study comprising 47 toddlers (*n* = 27 male) in an age range of 9 to 16 months (*M* = 12.18 months, *SD* = 2.11). Mean birth weight was 3482 g (SD = 488, range: 2190 – 4450 g) and the APGAR score was either 9 or 10 (*M* = 9.93, *SD* = 0.25) after 10 minutes after birth. All parents gave written informed consent. The study was approved by the ethics committee of the Institute of Psychology at the Humboldt University of Berlin, Germany.

### Sleep Recordings and Scoring

Sleep was recorded using a portable amplifier (LiveAmp, Brain Products GmbH, Gilching, Germany) with a sampling rate of 500 Hz. Channel Cz served as recording reference. EEG was recorded from 26 channels. Additionally, electromyogram (EMG) and vertical and horizontal electrooculogram (EOG) were recorded. Offline, the EEG was re-referenced to the average of the left and right mastoids (TP9, TP10) and filtered between 0.5 and 35 Hz. Sleep scoring was based on standard criteria (Berry et al., 2017; Grigg-Damberger et al., 2007; Scholle & Feldmann-Ulrich, 2012) using channels F3, F4, C3, C4, O1 and O2 as well as EOG and EMG signals. Sleep was visually scored by two independent raters with high consistency (> 90 %). For each nap, total sleep time and the time spent in sleep stages N1, N2, N3, and REM were determined.

Artefactual epochs (30 s epochs) were marked during scoring and excluded from further analyses. Furthermore, bad channels were visually identified and interpolated for all further analyses. In 21 participants at least one channel was interpolated (*M* = 1.76, *SD* = 1.18, range: 1-5).

### Spindle Detection

Sleep spindle detection was based on the individual frequency peaks in the fast spindle frequency range (12-16 Hz). Thus, for each participant power-spectra of all artefact-fee non-REM (N2 and N3) epochs were calculated and the individual peak frequency determined. Next, spindles were detected using the MATLAB-based toolbox SleepTrip (RRID:SCR_017318) which is based on Mölle et al. (2002) and has previously been used in adult and developmental studies (Bastian et al., 2022; Kurz et al., 2024).

For each individual, the signal was low-pass filtered at 30 Hz, down sampled (100 Hz) and then bandpass filtered at ±1.5 Hz around the identified frequency peak (FIR filter, −3dB at frequency peak ±2 Hz). Then, for each sample point, the root mean square (RMS) in a 0.2-second time window is calculated, which is further smoothed by an averaging time window of 0.2 seconds. If the smoothed moving RMS signal exceeds the filtered signal in the respective channel by 1.5 SDs over a period of 0.5 to 3 seconds, a spindle is detected. Two succeeding spindles were considered as one spindle if the interval between the end of the first spindle and the beginning of the second spindle was shorter than 0.5 seconds and the resulting merged spindle was not longer than 3 seconds.

### Slow Oscillation detection

SOs were similarly detected during non-REM sleep using the SleepTrip toolbox. The signal was first low-pass filtered at 30 Hz and down sampled (100 Hz). Then a DC removal procedure and a 3.5 Hz low-pass filter was applied (FIR filter, −3dB at 3.5 Hz). In the filtered signal, all intervals with consecutive positive-to-negative zero crossings with a duration of 0.75 to 2 seconds (corresponding to a frequency between 0.5 and 1.33 Hz) were marked as potential SOs. Then a SO was detected when the negative-to-positive peak amplitude was 1.25 times greater than the mean negative-to-positive peak amplitude of all potential SOs within the channel and the negative peak (trough) amplitude was 1.25 times lower than the mean negative peak amplitude of all potential SOs in this channel.

### Co-occurrence of SOs and spindles

In a first step, we looked for each detected spindle, whether its center (i.e., the maximum trough) occurred within the two positive-to-negative zero crossings of an SO. Co-occurrence rates were calculated as the percentage of spindles within a channel that co-occurred with an SO, further referred to as observed co-occurrence rates. In order to investigate whether observed co-occurrence rates differed from chance, we calculated for each channel the chance level in the same way as previously done in Kurz et al. (2024), which is the percentage of non-REM sleep with SOs (total SO duration/non-REM sleep duration × 100).

In a second step, the ± 1.5 s interval around the negative peak (trough) of all SOs was subdivided into 15 200 ms bins to calculate peri-event time histograms (PETHs; cf. Mölle et al., 2011; Muehlroth et al., 2019). For each bin, occurrences of spindle events (i.e., all troughs and peaks of the detected spindles) were summed across all SOs of a channel of an individual and then normalized by the number of spindle events co-occurring with all SOs in the respective channel.

Next, we calculated the time-frequency representations (TFRs) of SOs using the open-source toolbox FieldTrip (Oostenveld et al., 2011). For this, the ±3 seconds around the SO trough were extracted and subjected to time-frequency analyses using Morlet wavelets with linearly increasing cycles from 4 to 12, in a frequency range from 2 to 20 Hz in steps of 0.5 Hz. The power in each channel was normalized by the average power in the ±1.5 seconds around the SO trough, represented as percentage change, with positive values indicating an increase in power from the baseline.

Lastly, we extracted for each individual and channel the SO phase at the spindle amplitude maximum. Using the CircStat toolbox (Berens, 2009), we calculated the mean resultant vector length and the mean phase within each and across participants for each channel. The mean resultant vector length ranges from 0 to 1 and measures of how consistently maximum spindle amplitudes are clustered toward the same SO phase, with 1 indicating maximum clustering toward the same phase. We further calculated an upstate alignment score to investigate age-dependent changes of phase distributions. This phase score was calculated as the cosine of the angular distance between each SO phase at the spindle maximum and the SO upstate (0°). Higher values indicate that spindle maxima occurred closer to the SO upstate. A positive association with age would therefore indicate increasingly upstate-centered coupling.

Due to the differing and sometimes very low number of co-occurring events across participants, we performed sensitivity analyses for which we calculated surrogate data. For each participant and channel, we generated a null distribution by randomly sampling one time point within each SO that co-occurred with a spindle and extracting the SO phase at these random time points. This procedure was repeated 5000 times for each participant and channel. The surrogate-corrected resultant vector length was calculated as the difference between the observed resultant vector length and the mean surrogate resultant vector length, divided by the standard deviation of the surrogate distribution. Thus, this measure expresses phase consistency relative to an event-matched null distribution and accounts for the fact that nonzero vector lengths can arise by chance, particularly when the number of cooccurring events is small. Higher values indicate that spindle amplitude maxima were more consistently aligned to a specific SO phase than expected if their timing within the same cooccurring SOs were random.

Similarly, the surrogate corrected upstate alignment score was calculated as the difference between the observed phase score and the mean surrogate phase score, divided by the standard deviation of the surrogate distribution.

### Statistical Analysis

All statistical analyses were done in R version 4.6.1 (R Core Team, 2026) and MATLAB R2023b. Age was either treated continuously (age in days) or as a factor to compare participants younger (*n* = 28) and older (*n* = 19) than 12 months. To maintain comparability with our previous study (Kurz et al., 2024), we opted for a two-fold approach – a cluster-based approach to investigate whether age-dependent changes in coupling occur across a spatial cluster and an average-based approach, where we averaged across, frontal, central, parietal and occipital sites. Thus, depending on the analysis either all available channels were considered or the average of frontal (F3, Fz, F4), central (C3, Cz, C4), parietal (P3, Pz, P4) and occipital (O1, O2) electrodes was used.

Linear mixed effects models were calculated in R using the library lme4 (Bates et al., 2015) and lmerTest (Kuznetsova et al., 2017) to obtain F-statistics and *p*-values based on Type III sum of squares and Satterthwaite’s method. Simple slopes analysis and pairwise comparisons (Bonferroni corrected) were done using the emmeans package (Lenth, 2023). All models included a random intercept for each participant.

Age group comparisons (independent samples t-tests) or associations (Spearman correlation) between an EEG-based measure with age across all electrodes were done in MATLAB using the Fieldtrip toolbox (Oostenveld et al., 2011) and corrected for multiple comparisons using cluster-based permutation tests (Maris & Oostenveld, 2007) with each 1000 permutations.

Co-occurrence rates of spindles and SOs were analyzed by linear-mixed effects models. The first model included the fixed effects age group (< 12 months, > 12 months), channel (frontal, central, parietal, occipital) and ratio type (observed, chance-level). A second model analyzing only observed co-occurrence rates included the fixed effects age (continuously) and channel (frontal, central, parietal, occipital).

PETHs of the whole group and separately for participants younger and older than 12 months were tested against surrogate data (average of 1000 times bin shuffled data) for each channel using cluster-based permutation tests. Similarly, the PETHs of the age groups were compared at each channel and corrected for multiple comparisons using cluster-based permutation tests.

Similarly, baseline normalized TFRs of the whole group and separately of participants younger and older than one year were tested against zero using cluster-based permutation tests. Additionally, the baseline normalized TFRs of the two age groups were compared.

Using spearman correlations, TFRs across the whole sample were correlated with age and corrected for multiple comparisons using cluster-based permutation tests. Additionally, baseline normalized power in the spindle frequency range (12-16 Hz) around the SO upstate (200 to 900 ms after the SO trough) was extracted and averaged across frontal, central, parietal and occipital channels and subjected to a linear mixed-effects model including fixed effects for age and channel.

Rayleigh tests within and across participants were calculated using the CircStat toolbox (Berens, 2009) to test whether SO phases at spindle amplitude maxima were distributed (non)-uniformly. Mean resultant vector lengths and phase scores were analyzed using linear-mixed effects models, including the fixed effects channel (Fz, Cz) and age (continuously).

All tests were two-tailed and the critical alpha level at .05.

## Results

Investigation of spindle and SO characteristics did not show a main effect of age group (< 12 months compared to > 12 months), or an interaction between age group and channel (frontal, central, parietal, occipital) for any measure except the SO slope (age group × channel interaction: *F*(3,141) = 2.97, *p* = .034). SO slopes did not differ between the two age groups in any channel (all *p* > .999). In infants younger than 12 months, the occipital slope was steeper than in all other channels (all *p* < .002). In participants older than 12 months, there was only a difference between the parietal and occipital slope (*p* = .046). All other spindle and SO measures showed topographical effects (all *p* < .001). While spindles were denser at fronto-central sites (Figure 1A), SOs showed highest densities occipitally (Figure 1B). See Table S1 for means and standard errors of all spindle and SO characteristics.

**Figure 1.**
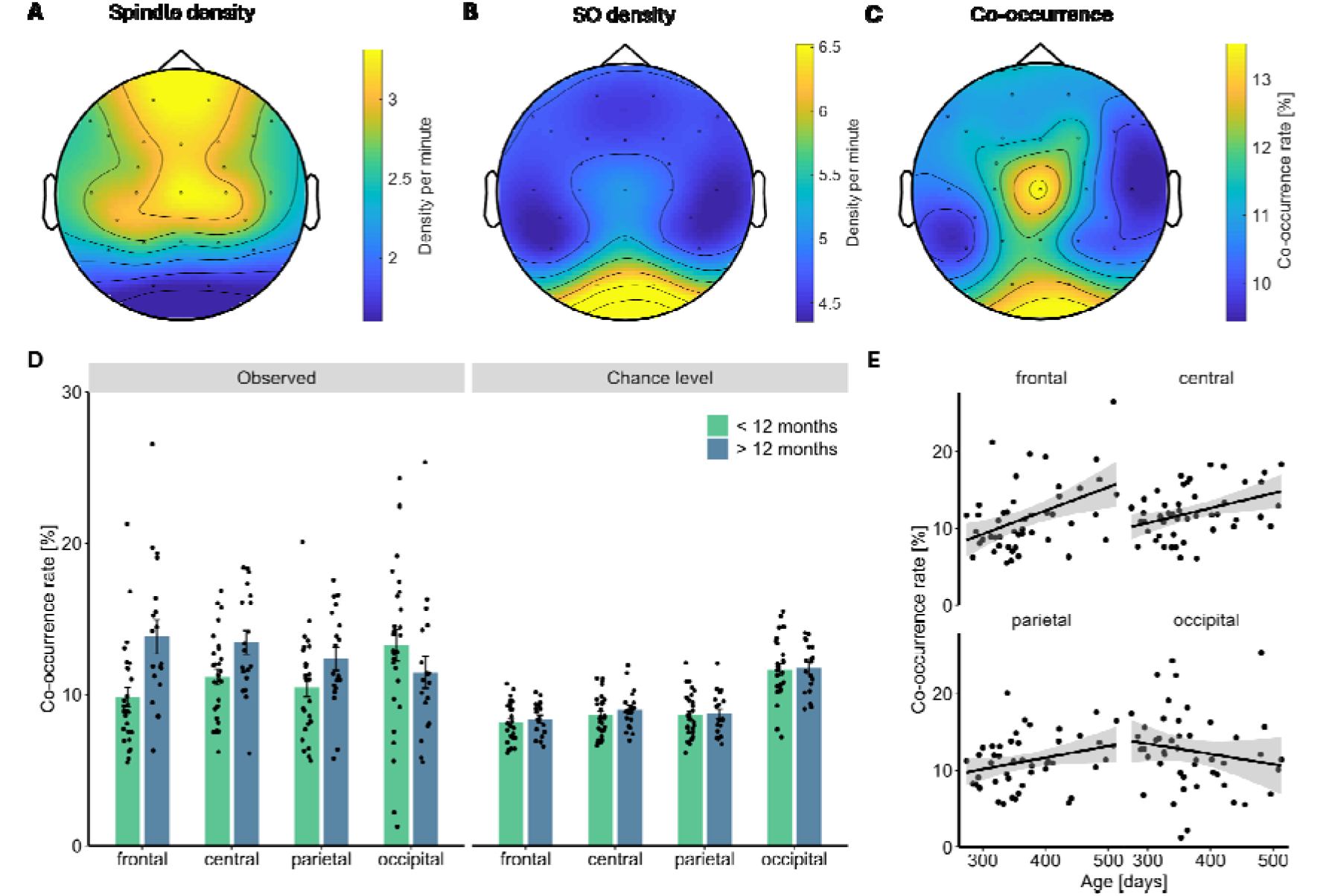
Spindle density (**A**), SO density (**B**), rate of spindles co-occurring with SOs (**C**). **D**: Cooccurrence rates in participants younger and older than 12 months, separately at the average of frontal, central, parietal and occipital channels, and observed and chance-level rate. **E**: Association between age and observed co-occurrence rate at the average of frontal, central, parietal and occipital channels. Shaded areas represent the 95% confidence interval. Please note the figure is illustrative, statistical inference is based on the linear mixed-effects model. See Figure S1 for co-occurrence rates in all channels.

Next, we examined the ratio of spindles co-occurring with SOs. In the overall group, co-occurrence rates were highest at channel Cz and occipital electrodes (Figure 1C). There was a significant three-way interaction between age group, channel and ratio type (*F*(3,329) = 5.06, *p* = .002, Figure 1D). In participants older than 12 months, co-occurrence rates had a mid-central maximum and exceeded chance-level at frontal (*p* < .001), central (*p* < .001) and parietal (*p* < .001), but not occipital sites (*p* > .999). Co-occurrence rates in infants younger than 12 months were highest in the occipital regions, and exceeded chance-level at central sites (*p* = .003) only (Figure S1, see Table S2 for means and standard errors). The two age groups did not differ in their chance-level estimates at any site (all *p* > .999), but observed co-occurrence rates were greater in participants older than one year compared to those younger than one year only frontally (*p* < .001). Although participants with steeper SO slopes had greater co-occurrence rates (*F*(1,120.02) = 10.88, *p* = 001), adding the SO slope as a covariate did not change the results.

Across all children, an association between observed co-occurrence rates and age (continuously) was seen in frontal (simple slopes: *t*(157) = 3.34, *p* = .001) and central (simple slopes: *t*(157) = 2.12, *p* = .036), but not parietal (*p* = .102) and occipital channels (*p* = .146, age × channel interaction: *F*(3,141) = 6.15, *p* < .001, Figure 1E). Frontal, central and parietal channels did not differ from each other in their simple slopes (all *p* > .941). Frontal and central channels had a greater age-dependent increase in observed co-occurrence rates than occipital channels (all *p* < .019). Controlling for the SO slope left the reported results unchanged.

In the next step we investigated peri-event time histograms, i.e. the SO time-binned spindle event occurrence rates. Across the whole group spindle event occurrence compared to surrogate data was increased at fronto-central regions around the SO upstate (cluster *p* < .001, see Figures S2 and 2A). Descriptively this pattern was less pronounced in individuals below one year (see Figure 2B for co-occurrence rates at exemplary channel Cz and Figures S3-S4 for results across all channels), but no significant cluster emerged when comparing the two age groups regarding the spatio-temporal distribution of spindle event occurrence rates. However, when exploring continuous age-related changes regarding the SO-bin specific spindle event rates at channel Cz, we found an increase with age at the time bins covering 100 to 700 ms, which includes the peak of the SO (Figure 2C). The bin with the highest correlation coefficient (i.e., in the range of 300 to 500 ms) differed significantly from all other correlation coefficients except that in the range of 100 to 300 ms (z = 1.52, p = .129).

**Figure 2.**
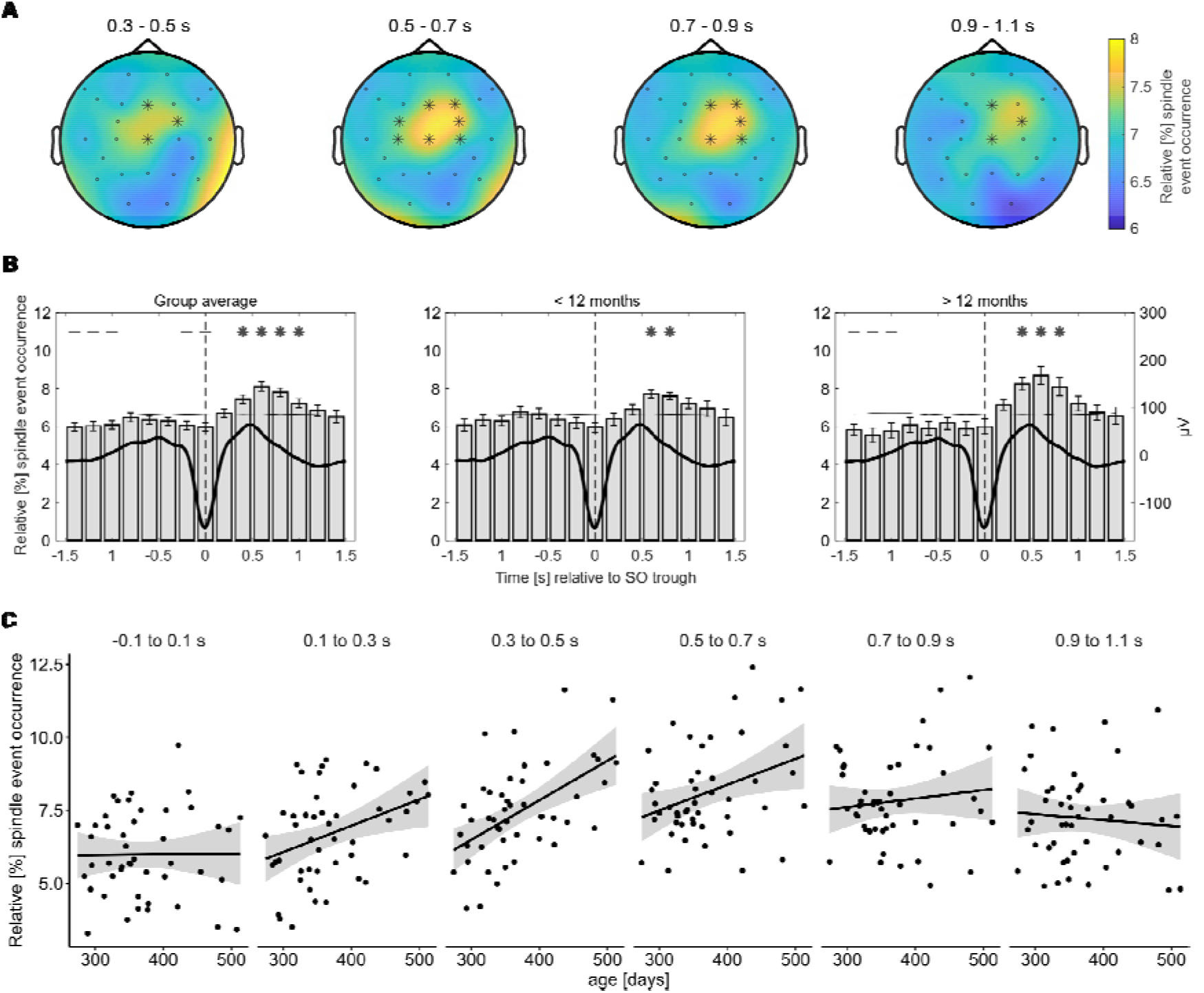
Peri-event time histograms. **A**: Tested against surrogate data, relative spindle rates (in %) increasingly occurred around the SO upstate across all participants at fronto-central sites. **B**: Relative spindle event occurrence at exemplary channel Cz, separately across the whole group (left), individuals younger than 12 months (middle) and older than 12 months (right). Asterisks depict significantly greater relative spindle occurrence compared to surrogate data. Dashed lines depict significantly lower relative spindle occurrence compared to surrogate data (thin horizontal line). See Figures S2-S4 for results across all channels. **C**: Correlations between age and the relative spindle event occurrence, separately for different time bins around the SO trough at channel Cz.

Next, we examined SO trough-locked power using time frequency representations. Across all participants, we observed one big cluster spanning across all channels (cluster *p* < .001, Figure S4). At the SO trough all channels showed an increase in activity in a wide range of frequencies (mostly ranging from 4 to ~11 Hz and ~15 to 20 Hz) compared to baseline. Additionally, activity was greater compared to baseline in the spindle frequency range (~ 10 to 16 Hz) during the SO upstate (~250 to 900ms) at fronto-central channels. A similar pattern of results was seen in both participants younger and older than 12 months with each again one big positive cluster (cluster *p* < .001). See Figures S5 and S6 for results in all channels and Figure 3A and 3B for exemplary channel Cz. Nonetheless, power change from baseline was greater in participants older than 12 months compared to those younger than 12 months mainly in fronto-central areas (cluster *p* = .015, see Figure 3C for exemplary channel Cz and Figure S7 for all channels). Accordingly, spindle activity around the SO upstate was associated with age across the whole group (cluster *p* = .004, see Figure 3D for the association at exemplary channel Cz and Figure S8 for the result across all channels). To test whether the age-group difference in SO trough-locked spindle power was explained by age-related differences in SO morphology, we repeated the TFR group comparison after regressing out the channel-specific SO slope from the TFR power values at each channel, frequency, and time point. The resulting residualized TFRs were submitted to the same cluster-based permutation test. This sensitivity analysis yielded the same group differences as described above.

**Figure 3.**
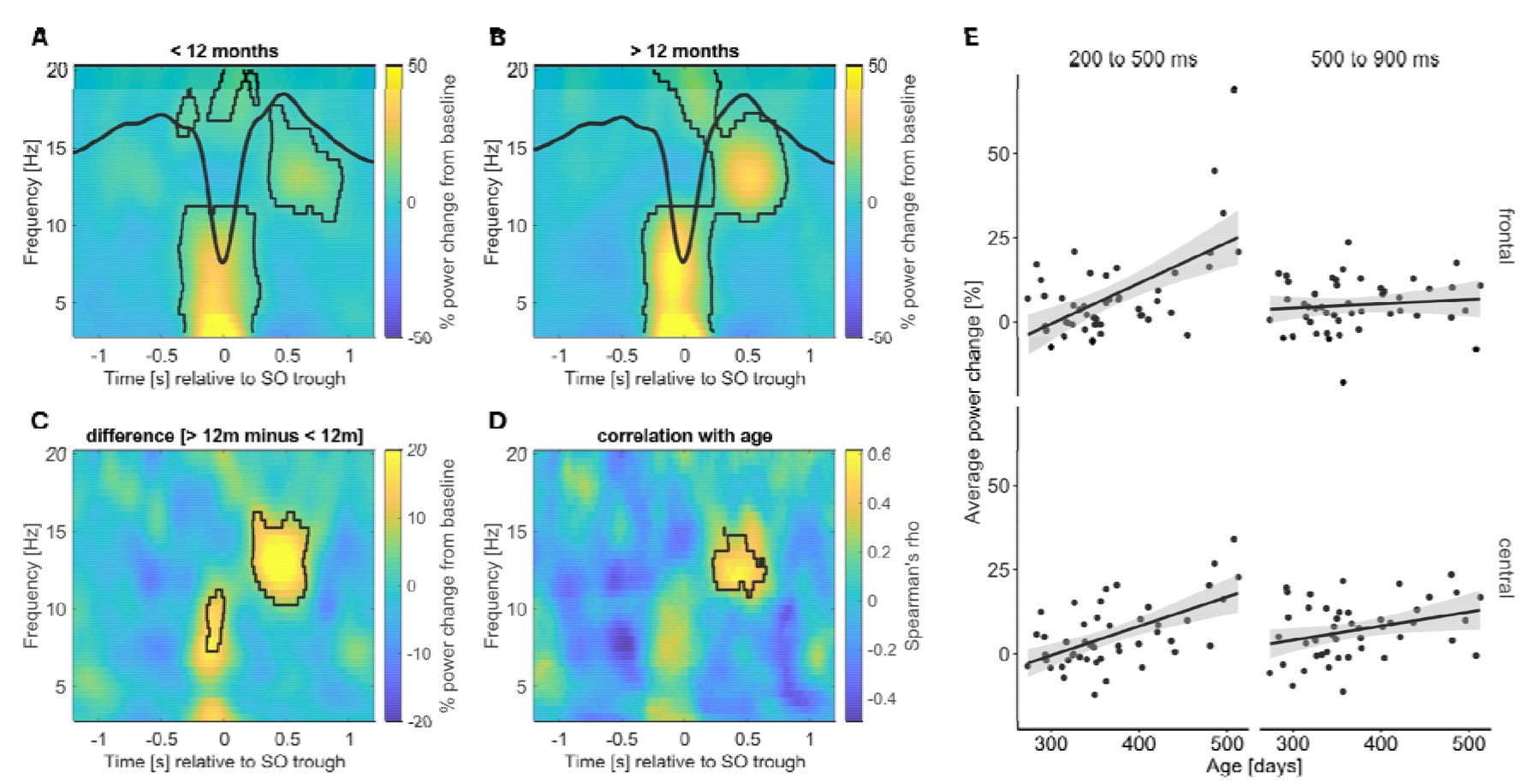
Time-frequency representation of SO events at exemplary channel Cz in participants younger than 12 months (**A**) and older than 12 months (**B**), as well as the difference between the age groups (**C**). Plots depict the change from baseline (in %). **D**: Correlation of time-frequency representation of SO events with age in days at exemplary channel Cz. Black lines indicate the outline of significant positive clusters. See Figures S5 – S8 for results across all channels. **E**: Association between age and the average spindle activity (12-16 Hz) during the SO upstate (separately for 200-500 ms and 500-900 ms after the SO trough). Please note Figure E is illustrative, statistical inference is based on the linear mixed-effects model.

In order to exploratively test whether this increase in spindle power around the SO upstate varies temporally with age, we subjected the average baseline-normalized spindle power (12-16 Hz) between 200 to 500 ms and 500 to 900 ms after the SO trough at the average of frontal, central, parietal and occipital channels to a linear mixed-effects model. This revealed a significant interaction between age (continuous), time and channel (*F*(3,329) = 5.23, *p* = 002). Simple slopes analyses showed an age-dependent increase in the early time window at frontal (*t*(388) = 7.55, *p* < .001) and both time windows at central channels (200 to 500 ms: *t*(388) = 5.43, *p* < .001, 500 to 900 ms: *t*(388) = 2.61, *p* = .009, Figure 3E) but at no time window parietally or occipitally (all *p* > .142). The frontal age-slope was steeper in the early than the late time window (*t*(344) = 4.88, *p* < .001). Furthermore, age-slopes did not differ between frontal and central channels in the early and late time window (all *p* > .999). In the early time window, age-slopes of frontal and central channels were each greater than in occipital channels (all *p* < .001) and the frontal age-slope was greater than the parietal age-slope in the early time window (*p* < .001). Controlling for the SO slope did not change the results and the SO slope was not associated with the averaged spindle power (*F*(1,72.02) = 1.35, *p* = .249).

Paralleling these findings, age groups differed at frontal and central channels at 200-500 ms (all *p* < .001) but not 500-900 ms after the SO trough (all *p* > .278), although here the three-way interaction did not reach significance (age group × time × channel: *F*(3,329) = 2.06, *p* = .106). Analysis of SO peak- instead of SO trough-locked TFRs showed that both age groups had increased activity in the spindle frequency range around the SO peak (Figures S9-S11) and confirmed that participants older compared to younger than 12 months had greater spindle power mainly before the upstate peak (cluster *p* = .016, Figure S9C, S12).

Finally, we investigated SO-spindle coupling defined as the SO phase at the spindle amplitude maximum (Figure 4). Only channels with at least five co-occurring events were considered. Rayleigh tests at the individual participant and channel level indicated that SO phases at spindle amplitude maxima were mostly uniformly distributed. The channel with the highest proportion of non-uniform phase distributions was channel Cz, where 12 out of 45 participants had a non-uniform distribution. Due to the differing number of participants providing data for a specific channel and the former analyses revealing no temporally coordinated coalescence between spindles and SOs at parietal and occipital sites, we opted for only including channels Fz and Cz in the following analyses. Rayleigh tests across participants revealed uniform distributions of mean phases at channel Fz (Rayleigh-Z = 2.16, *p* = .116, Figure 4A) and a non-uniform distribution at Cz (Rayleigh-Z = 9.76, *p* < .001, Figure 4B). All following models investigating age-dependent changes in phase consistency and precision included the covariates number of co-occurring events and the SO slope.

**Figure 4.**
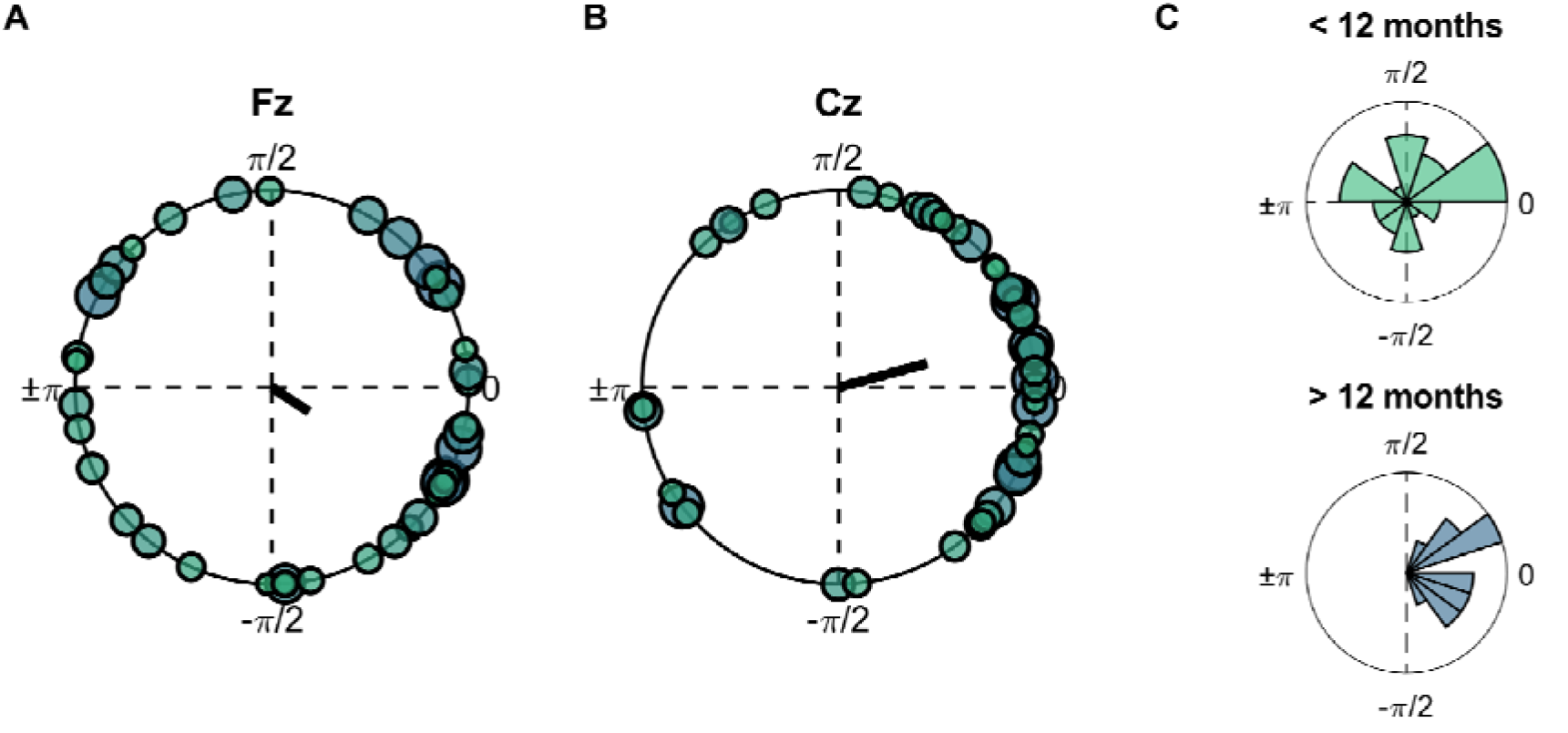
Mean phases across participants at channel Fz (**A**) and Cz (**B**). The direction of the vector depicts the circular group mean, with its length depicting the consistency of coupling within the channel. Larger circles indicate older participants. **C:** Two exemplary polar-histograms of one participant younger (upper panel) and one participant older (lower panel) than 12 months.

First, we investigated the mean resultant vector length, i.e. how consistently spindle maxima occurred at the same SO phase. There was no association between the mean resultant vector length and age (*F*(1,44.03) = 2.66, *p* = .110) and no interaction between age and channel (*F*(1,44.36) = 0.93, *p* = .339). Participants with a lower number of co-occurring events (*F*(1,53.09) = 14.86, *p* < .001) and steeper SO slopes (*F*(1,56.71) = 7.76, *p* = .007) showed higher phase consistency. We then performed sensitivity analyses to examine whether this null result was influenced by unreliable vector-length estimates in participants with few co-occurring events. First, we repeated the analysis in participants with at least 10 co-occurring events. A significant age and channel interaction (*F*(1,35.00) = 4.45, *p* = .042) showed that phase consistency increased with age at channel Cz (simple slopes: *t*(73.6) = 2.70, *p* = .009) but not at Fz (*p* = .960). Age trends differed between channel Cz and Fz (*p* = .0499). Participants with less co-occurring events (*p* = .007) and a steeper slope (*p* = .027) had more consistent coupling towards the same SO phase.

Because this event-count threshold further reduced the sample size, we additionally computed a surrogate-corrected measure of phase consistency using the full sample with at least 5 co-occurring events (see methods). Using this surrogate-corrected vector length, there was no interaction between age and channel (*p* = .337), but age was associated with increased phase consistency across channels (*F*(1,44.59) = 6.21, *p* = .016). The number of co-occurring events was not associated with phase consistency (*p* = .979), but a steeper SO slope was still associated with greater phase consistency (*p* = .019).

Lastly, we examined whether the phases are closer to the SO upstate in older participants to investigate whether coupling precision increases with age. Using an upstate alignment score, we started again with the threshold of at least five co-occurring events. There was a trend that the mean phase was more centered to the SO upstate with increasing age (*F*(1,43.38) = 3.38, *p* = .073). Steeper slopes and a lower number of co-occurring events were associated with greater upstate centered coupling (*p* < .001 and *p* = .053, respectively). Using the threshold of at least 10 co-occurring events, strengthened the age-dependent increase in phase precision (*F*(1,31.61) = 4.78, *p* = .036). Using the surrogate-corrected phase score similarly revealed more upstate centered coupling with increasing age across channels (*F*(1,44.38) = 6.83, *p* = .012) even when using the participants with at least 5 cooccurring events. A steeper SO slope (*p* < .001) but not the number of co-occurring events (*p* = .358) was associated with increased phase precision. The same pattern of results emerged when using the absolute distance to the SO upstate instead of the cosine of the distance between the SO phase at maximum spindle amplitude and SO upstate.

## Discussion

We here studied the early development of the coordinated coupling of spindles and SOs, a mechanism that has repeatedly been demonstrated to support memory consolidation in children and adults (Hahn et al., 2020; Kurz et al., 2023; Muehlroth et al., 2019; Ng et al., 2025). We found that the nesting of spindles into the upstate of SOs emerges already in the first year of life over the fronto-central cortex and increases in strength and precision with age. The frequent co-occurrence of spindles and SOs at occipital regions in infants did not exceed chance level and was also not temporally coordinated. Together, these findings indicate that the early development of SO–spindle coupling is characterized by the progressive refinement of fronto-central temporal coordination.

Across early life, the characteristics and the topographical predominance of SOs undergo pronounced developmental changes (Kurth et al., 2010; Page et al., 2020), which has been proposed to reflect ongoing cortical maturation and synaptic remodeling (Buchmann et al., 2011; Fattinger et al., 2014; Feinberg & Campbell, 2010). Similarly, spindle characteristics are subject to developmental changes (Kwon et al., 2023). Investigating a relatively narrow age range of eight months, here, we found no differences in the characteristics of spindles and SOs between participants younger and older than one year, except for the SO slope. Although infants and toddlers did not differ in SO slope at any region, infants showed steepest SO slopes occipitally, while toddlers showed overall smaller differences between cortex regions. This occipital predominance in infants is consistent with previous observations of particularly steep slow-wave slopes over occipital cortex during the first year of life (Fattinger et al., 2014). We thus added the SO slope as covariate to the statistical models investigating age-dependent changes in the temporal coordination of SO and spindles. Steeper slopes were associated with various measures such as increased cooccurrence rates and closer coupling of spindles to the upstate of SOs, suggesting that a steeper slope contributes to a temporally coordinated coupling. However, none of the age-dependent changes in coupling measures were affected by including the SO slope in the analysis. This means that the morphology of SOs contributes to interindividual variability in SO-spindle coupling but does not account for its age-related development. Instead, the increasing temporal precision of SO–spindle coupling may reflect a growing maturity of the mechanisms involved in the temporal coordination between the underlying oscillatory processes.

Including participants from 9 to 16 months, here we extend the age range of our previous study that compared SOs, spindles and their coupling in 2 to 3 months old infants with those of 14 to 17 months old toddlers (Kurz et al., 2024). In line with our previous study, co-occurrence rates of spindles with SOs were higher in toddlers than infants, but this increase was limited to frontal cortex regions for the adjacent age groups compared here. In the participants younger than one year, co-occurrence rates of spindles with SOs reached their maximum in the occipital cortex, which corresponds with the findings in 6-month-old infants (Jaramillo et al., 2023). However, by analyzing what is expected by chance, we could specify that the co-occurrence rates in infants exceeded chance level only at the central region. Thus, the observed high co-occurrence rates in the occipital cortex in infants appears to reflect the posterior predominance of slow-wave activity (Beaugrand et al., 2026; Fattinger et al., 2014), rather than a temporal coordination of spindles with SOs at occipital sites. Since the predominance of slow-wave activity shifts from posterior to anterior regions across development (Kurth et al., 2010), we recommend to include an analysis of the chance-level of co-occurrence as a general practice, when reporting observed co-occurrence rates of spindles and SOs in children.

Besides this rather coarse measure of the temporal coordination between sleep spindles and SOs, we used several measures to investigate the coupling of spindles to the upstate of SOs and its developmental changes in the very early life. These included the analyses of PETHs with spindle event occurrence rates binned into 200 ms in the 1.5 seconds around the SO trough, TFRs with power changes from baseline during SO events, and SO phases at spindle amplitude maxima. Although these individual measures were based on slightly different criteria (e.g., time-windows around the SO trough, restriction to coupled vs. uncoupled events), they all yield a similar picture: The temporally coordinated coupling of spindles to the SO upstate emerges in the fronto-central cortex, where it increases with age, and is absent at parietal and occipital regions. Moreover, age-related increases in co-occurrence rates and spindle activity before and after the SO upstate did not differ between frontal and central regions. Both, correlation of individual bins in PETHs and analysis of time-frequency representations revealed that age-related changes in spindle activity are most prominent right before the SO upstate peak. In conjunction with the agerelated shift of spindle maxima toward the upstate peak, this reflects a developmental refinement in the timing of spindle activity relative to the SO peak rather than a broad increase in spindle activity across the entire SO upstate.

Restricting the analysis to events that are coupled within the two positive-to-negative zero crossings of an SO, as was done for extracting the SO phase at the spindle amplitude maxima, resulted in a low number of contributing events for some participants. This is particularly relevant because estimates of phase consistency based on the mean resultant vector length are sensitive to the number of contributing events, especially at small sample sizes (Andrzejak et al., 2023). Implementing several sensitivity analyses for phase consistency as well as phase angle, suggested that age-related changes in preferred SO phase may only be detectable when phase estimates are based on a sufficient number of cooccurring events. Low event counts may obscure developmental changes in both preferred phase and phase consistency. Our analyses suggested that phase consistency tends to increase with age, although it remains unclear whether this is the case for midcentral regions only or also for midfrontal areas. Furthermore, the older the participant the closer were spindle amplitude maxima coupled to the SO upstate. This finding was consistent across our different sensitivity analysis. Overnight sleep recordings or pooling data of multiple naps in the same infants or toddlers could serve as alternative approaches to obtain a sample with a sufficient number of co-occurring events and replicate our findings without surrogate-corrected measures.

In conclusion, our findings suggest that the temporally coordinated coupling of SOs and spindles first emerges in fronto-central cortex regions in the second half of the first year of life and increases gradually in precision during the subsequent months of life. Importantly, high rates of SO–spindle co-occurrence alone did not necessarily indicate the presence of coupling, emphasizing the relevance of considering both chance-level co-occurrence and the precise temporal relationship between the two oscillations. Together, our findings identify the transition from infancy to toddlerhood as an important period for the maturation of SO– spindle coupling. They thus provide a basis for investigating the functional significance of this coupling mechanism for memory consolidation in the earliest stages of life.

## Acknowledgements

We thank all families who participated in this study. Special thanks to Jördis Haselow, Katja Friedrich de Guzman, and Kirsten Bräuner for carrying out the experimental sessions. The study was supported by grants from the Deutsche Forschungsgemeinschaft to M.F. (468645090) and J.B. (FOR 5434).

## Data Availability

The data underlying this article will be shared on request to the corresponding author.

## References

Andrzejak, R. G., Espinoso, A., García-Portugués, E., Pewsey, A., Epifanio, J., Leguia, M. G., & Schindler, K. (2023). High expectations on phase locking: Better quantifying the concentration of circular data. Chaos: An Interdisciplinary Journal of Nonlinear Science, 33(9), 091106. 10.1063/5.0166468

Bastian, L., Kurz, E.-M., Näher, T., Zinke, K., Friedrich, M., & Born, J. (2024). Long-term memory formation for voices during sleep in three-month-old infants. Neurobiology of Learning and Memory, 215, 107987. 10.1016/j.nlm.2024.107987

Bastian, L., Samanta, A., Ribeiro de Paula, D., Weber, F. D., Schoenfeld, R., Dresler, M., & Genzel, L. (2022). Spindle–slow oscillation coupling correlates with memory performance and connectivity changes in a hippocampal network after sleep. Human Brain Mapping, 43(13), 3923–3943. 10.1002/hbm.25893

Bates, D., Mächler, M., Bolker, B., & Walker, S. (2015). Fitting linear mixed-effects models using lme4. Journal of Statistical Software, 67, 1–48. 10.18637/jss.v067.i01

Beaugrand, M., Jaramillo, V., Mühlematter, C., Schoch, S. F., Reicher, V., Markovic, A., & Kurth, S. (2026). Tracing infant sleep neurophysiology longitudinally from 3 to 6 months: EEG insights into brain development. npj Biological Timing and Sleep, 3(1), 9. 10.1038/s44323-026-00071-7

Berens, P. (2009). CircStat: A MATLAB Toolbox for Circular Statistics. Journal of Statistical Software, 31(10), 21. 10.18637/jss.v031.i10

Berry, R. B., Brooks, R., Gamaldo, C., Harding, S. M., Lloyd, R. M., Quan, S. F., Troester, M. M., Bradley, V., & Medicine, f. t. A. A. o. S. (2017). The AASM manual for the scoring of sleep and associated events: Rules, Terminology and Technical Specification (Vol. 2.4). American Academy of Sleep Medicine.

Brodt, S., Inostroza, M., Niethard, N., & Born, J. (2023). Sleep-A brain-state serving systems memory consolidation. Neuron, 111(7), 1050–1075. 10.1016/j.neuron.2023.03.005

Buchmann, A., Ringli, M., Kurth, S., Schaerer, M., Geiger, A., Jenni, O. G., & Huber, R. (2011). EEG Sleep Slow-Wave Activity as a Mirror of Cortical Maturation. Cerebral Cortex, 21(3), 607–615. 10.1093/cercor/bhq129

Fattinger, S., Jenni, O. G., Schmitt, B., Achermann, P., & Huber, R. (2014). Overnight Changes in the Slope of Sleep Slow Waves during Infancy. Sleep, 37(2), 245–253. 10.5665/sleep.3390

Feinberg, I., & Campbell, I. G. (2010). Sleep EEG changes during adolescence: an index of a fundamental brain reorganization. Brain and Cognition, 72(1), 56–65. 10.1016/j.bandc.2009.09.008

Friedrich, M., Molle, M., Born, J., & Friederici, A. D. (2022). Memory for nonadjacent dependencies in the first year of life and its relation to sleep. Nat Commun, 13(1), 7896. 10.1038/s41467-022-35558-x

Friedrich, M., Molle, M., Friederici, A. D., & Born, J. (2019). The reciprocal relation between sleep and memory in infancy: Memory-dependent adjustment of sleep spindles and spindle-dependent improvement of memories. Dev Sci, 22(2), e12743. 10.1111/desc.12743

Friedrich, M., Molle, M., Friederici, A. D., & Born, J. (2020). Sleep-dependent memory consolidation in infants protects new episodic memories from existing semantic memories. Nat Commun, 11(1), 1298. 10.1038/s41467-020-14850-8

Friedrich, M., Wilhelm, I., Born, J., & Friederici, A. D. (2015). Generalization of word meanings during infant sleep. Nat Commun, 6, 6004. 10.1038/ncomms7004

Galland, B. C., Taylor, B. J., Elder, D. E., & Herbison, P. (2012). Normal sleep patterns in infants and children: A systematic review of observational studies. Sleep Medicine Reviews, 16(3), 213–222. 10.1016/j.smrv.2011.06.001

Grigg-Damberger, M., Gozal, D., Marcus, C. L., Quan, S. F., Rosen, C. L., Chervin, R. D., Wise, M., Picchietti, D. L., Sheldon, S. H., & Iber, C. (2007). The visual scoring of sleep and arousal in infants and children. J Clin Sleep Med, 3(2), 201–240. 10.5664/jcsm.26819

Hahn, M., Joechner, A. K., Roell, J., Schabus, M., Heib, D. P., Gruber, G., Peigneux, P., & Hoedlmoser, K. (2019). Developmental changes of sleep spindles and their impact on sleep-dependent memory consolidation and general cognitive abilities: A longitudinal approach. Dev Sci, 22(1), e12706. 10.1111/desc.12706

Hahn, M. A., Heib, D., Schabus, M., Hoedlmoser, K., & Helfrich, R. F. (2020). Slow oscillation-spindle coupling predicts enhanced memory formation from childhood to adolescence. eLife, 9, e53730. 10.7554/eLife.53730

Jaramillo, V., Schoch, S. F., Markovic, A., Kohler, M., Huber, R., Lustenberger, C., & Kurth, S. (2023). An infant sleep electroencephalographic marker of thalamocortical connectivity predicts behavioral outcome in late infancy. NeuroImage, 269, 119924. 10.1016/j.neuroimage.2023.119924

Joechner, A.-K., Hahn, M. A., Gruber, G., Hoedlmoser, K., & Werkle-Bergner, M. (2023). Sleep spindle maturity promotes slow oscillation-spindle coupling across child and adolescent development. eLife, 12, e83565. 10.7554/eLife.83565

Klinzing, J. G., Niethard, N., & Born, J. (2019). Mechanisms of systems memory consolidation during sleep. Nature Neuroscience, 22(10), 1598–1610. 10.1038/s41593-019-0467-3

Kurdziel, L. B. F. (2019). The Memory Function of Sleep Across the Life Span. In S. K. Jha & V. M. Jha (Eds.), Sleep, Memory and Synaptic Plasticity (pp. 1–39). Springer Singapore. 10.1007/978-981-13-2814-5_1

Kurth, S., Ringli, M., Geiger, A., LeBourgeois, M., Jenni, O. G., & Huber, R. (2010). Mapping of cortical activity in the first two decades of life: a high-density sleep electroencephalogram study. Journal of Neuroscience, 30(40), 13211–13219. 10.1523/JNEUROSCI.2532-10.2010

Kurz, E. M., Bastian, L., Molle, M., Born, J., & Friedrich, M. (2024). Development of slow oscillation-spindle coupling from infancy to toddlerhood. Sleep Adv, 5(1), zpae084. 10.1093/sleepadvances/zpae084

Kurz, E. M., Zinke, K., & Born, J. (2023). Sleep electroencephalogram (EEG) oscillations and associated memory processing during childhood and early adolescence. Dev Psychol, 59(2), 297–311. 10.1037/dev0001487

Kuznetsova, A., Brockhoff, P. B., & Christensen, R. H. (2017). lmerTest package: tests in linear mixed effects models. Journal of Statistical Software, 82(13), 1–26.

Kwon, H., Walsh, K. G., Berja, E. D., Manoach, D. S., Eden, U. T., Kramer, M. A., & Chu, C. J. (2023). Sleep spindles in the healthy brain from birth through 18 years. Sleep, 46(4), zsad017. 10.1093/sleep/zsad017

Lenth, R. V. (2023). emmeans: Estimated Marginal Means, aka Least-Squares Means. In (Version R package version 1.9.0) https://CRAN.R-project.org/package=emmeans

Maris, E., & Oostenveld, R. (2007). Nonparametric statistical testing of EEG- and MEG-data. J Neurosci Methods, 164(1), 177–190. 10.1016/j.jneumeth.2007.03.024

Mölle, M., Bergmann, T. O., Marshall, L., & Born, J. (2011). Fast and slow spindles during the sleep slow oscillation: disparate coalescence and engagement in memory processing. Sleep, 34(10), 1411–1421. 10.5665/SLEEP.1290

Mölle, M., Marshall, L., Gais, S., & Born, J. (2002). Grouping of spindle activity during slow oscillations in human non-rapid eye movement sleep. J Neurosci, 22(24), 10941–10947. 10.1523/JNEUROSCI.22-24-10941.2002

Muehlroth, B. E., Sander, M. C., Fandakova, Y., Grandy, T. H., Rasch, B., Shing, Y. L., & Werkle-Bergner, M. (2019). Precise Slow Oscillation-Spindle Coupling Promotes Memory Consolidation in Younger and Older Adults. Scientific Reports, 9(1), 1940. 10.1038/s41598-018-36557-z

Ng, T., Noh, E., & Spencer, R. M. C. (2025). Bayesian meta-analysis reveals the mechanistic role of slow oscillation-spindle coupling in sleep-dependent memory consolidation. eLife, 13, RP101992. 10.7554/eLife.101992

Nir, Y., Staba, Richard J., Andrillon, T., Vyazovskiy, Vladyslav V., Cirelli, C., Fried, I., & Tononi, G. (2011). Regional Slow Waves and Spindles in Human Sleep. Neuron, 70(1), 153–169. 10.1016/j.neuron.2011.02.043

Oostenveld, R., Fries, P., Maris, E., & Schoffelen, J. M. (2011). FieldTrip: Open source software for advanced analysis of MEG, EEG, and invasive electrophysiological data. Comput Intell Neurosci, 2011, 156869. 10.1155/2011/156869

Page, J., Lustenberger, C., & Fröhlich, F. (2020). Nonrapid eye movement sleep and risk for autism spectrum disorder in early development: A topographical electroencephalogram pilot study. Brain Behav, 10(3), e01557. 10.1002/brb3.1557

Page, J., Lustenberger, C., & Fr□hlich, F. (2018). Social, motor, and cognitive development through the lens of sleep network dynamics in infants and toddlers between 12 and 30 months of age. Sleep, 41(4), zsy024. 10.1093/sleep/zsy024

R Core Team (2026). R: A Language and Environment for Statistical Computing. In https://www.R-project.org

Scholle, S., & Feldmann-Ulrich, E. (2012). Polysomnographic atlas of sleep-wake states during development from infancy to adolescence. Ecomed Medizin.

Staresina, B. P., Bergmann, T. O., Bonnefond, M., van der Meij, R., Jensen, O., Deuker, L., Elger, C. E., Axmacher, N., & Fell, J. (2015). Hierarchical nesting of slow oscillations, spindles and ripples in the human hippocampus during sleep. Nature Neuroscience, 18(11), 1679–1686. 10.1038/nn.4119

Steriade, M., Timofeev, I., & Grenier, F. (2001). Natural Waking and Sleep States: A View From Inside Neocortical Neurons. Journal of Neurophysiology, 85(5), 1969–1985. 10.1152/jn.2001.85.5.1969

