## Supporting Information for "From co-occurrence to increasingly precise temporal coordination: Development of slow oscillation–spindle coupling from infancy to toddlerhood"

**Table S1.** Means and standard errors (SEM) of spindle and SO characteristics

| | | < 12 months n = 28 | | > 12 months n = 19 | |
|---|---|---|---|---|---|
| | | M | SEM | M | SEM |
| spindle density [events per minute] | | | | | |
| | frontal | 3.00 | 0.09 | 3.14 | 0.14 |
| | central | 3.13 | 0.08 | 3.21 | 0.12 |
| | parietal | 2.62 | 0.12 | 2.55 | 0.19 |
| | occipital | 1.64 | 0.09 | 1.57 | 0.10 |
| spindle frequency [Hz] | | | | | |
| | frontal | 13.36 | 0.15 | 13.32 | 0.23 |
| | central | 13.46 | 0.14 | 13.43 | 0.23 |
| | parietal | 13.43 | 0.15 | 13.42 | 0.26 |
| | occipital | 13.17 | 0.15 | 13.21 | 0.24 |
| spindle amplitude [µV] | | | | | |
| | frontal | 39.94 | 2.10 | 40.19 | 2.22 |
| | central | 38.10 | 1.98 | 33.87 | 1.54 |
| | parietal | 23.93 | 1.26 | 20.82 | 0.74 |
| | occipital | 20.40 | 1.01 | 17.91 | 0.98 |
| SO density [events per minute] | | | | | |
| | frontal | 4.60 | 0.11 | 4.73 | 0.11 |
| | central | 4.86 | 0.12 | 5.01 | 0.12 |
| | parietal | 4.96 | 0.14 | 5.03 | 0.17 |
| | occipital | 6.42 | 0.21 | 6.44 | 0.18 |
| SO amplitude [µV] | | | | | |
| | frontal | 246 | 9.93 | 233 | 10.24 |
| | central | 246 | 10.58 | 238 | 12.25 |
| | parietal | 216 | 9.91 | 209 | 11.96 |
| | occipital | 278 | 14.09 | 251 | 14.35 |
| SO slope [µV/s] | | | | | |
| | frontal | 640 | 26.03 | 618 | 29.68 |
| | central | 651 | 29.21 | 644 | 39.76 |
| | parietal | 585 | 27.38 | 574 | 33.45 |
| | occipital | 729 | 36.52 | 645 | 37.06 |
| SO duration [s] | | | | | |
| | frontal | 1.09 | 0.01 | 1.09 | 0.01 |
| | central | 1.09 | 0.01 | 1.10 | 0.02 |
| | parietal | 1.07 | 0.01 | 1.07 | 0.01 |
| | occipital | 1.11 | 0.01 | 1.12 | 0.01 |

**Table S2.** Co-occurrence rates [%] of spindles with slow oscillations

| | | < 12 months n = 28 | | > 12 months n = 19 | |
|---|---|---|---|---|---|
| | | M | SEM | M | SEM |
| observed | frontal | 9.81 | 0.65 | 13.81 | 1.12 |
| | central | 11.15 | 0.54 | 13.45 | 0.79 |
| | parietal | 10.45 | 0.61 | 12.36 | 0.76 |
| | occipital | 13.27 | 1.05 | 11.46 | 1.04 |
| chance | frontal | 8.18 | 0.25 | 8.37 | 0.25 |
| | central | 8.64 | 0.26 | 8.98 | 0.30 |
| | parietal | 8.62 | 0.28 | 8.71 | 0.32 |
| | occipital | 11.63 | 0.42 | 11.75 | 0.38 |

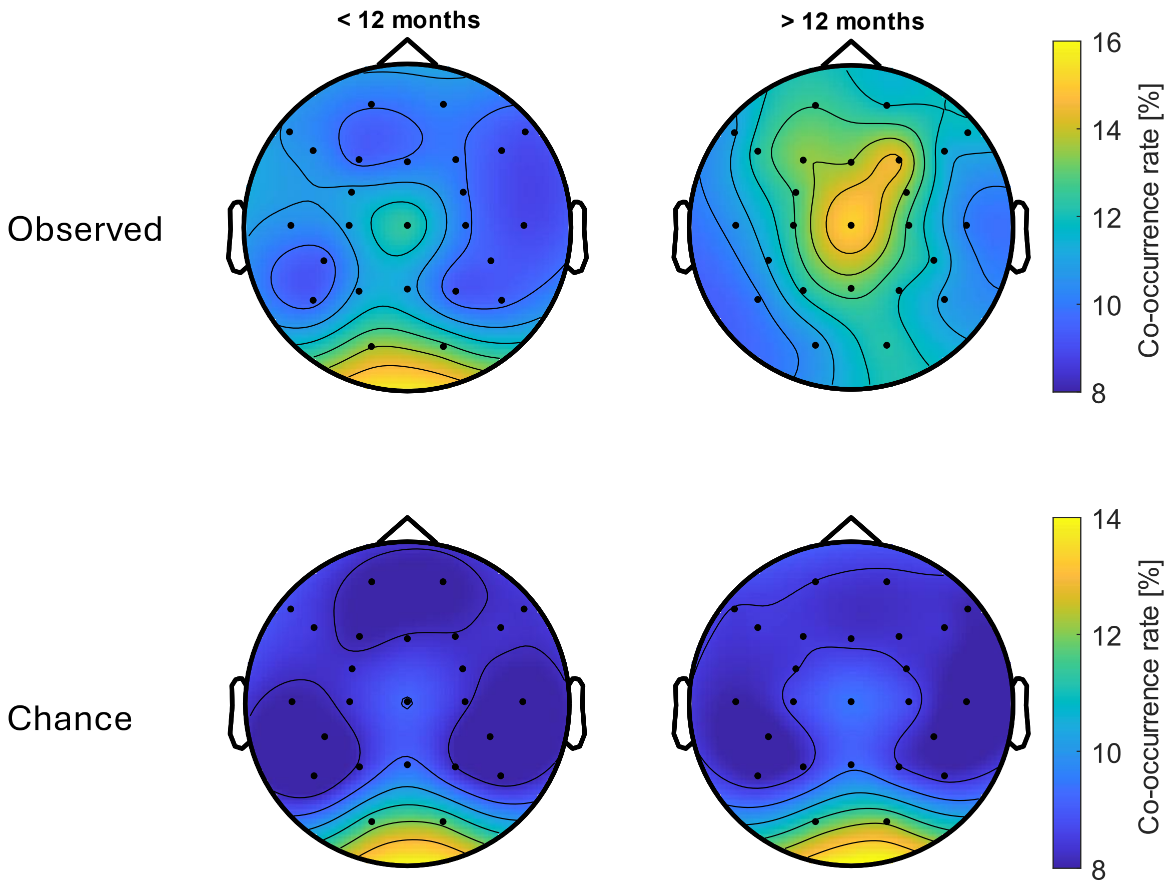


**Figure S1.** Co-occurrence rates of spindles with SOs, separately for observed co-occurrence rates, and chance-level rates.

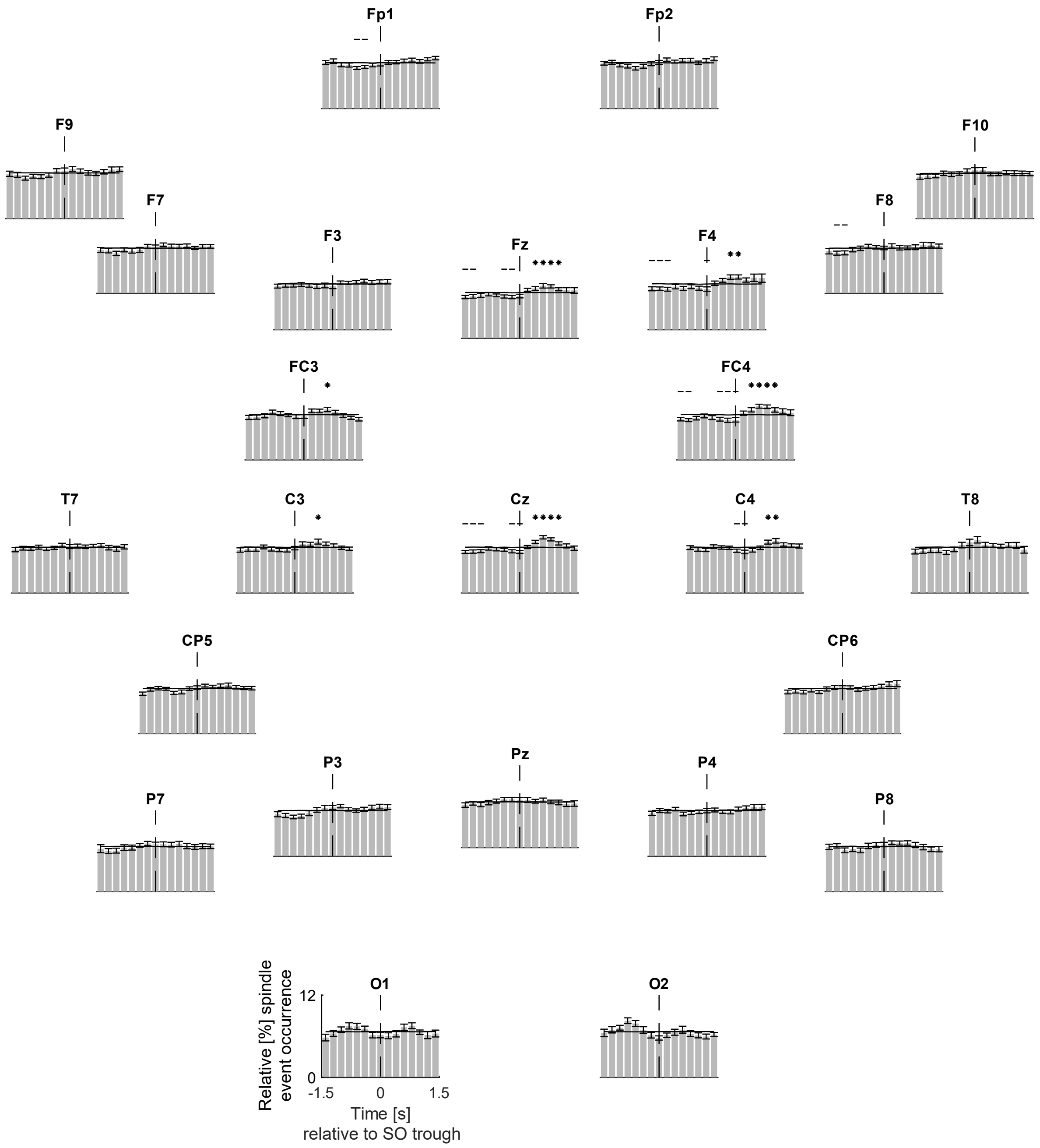


**Figure S2**. Peri-event time histograms across all participants. Asterisks depict significantly greater relative spindle occurrence compared to surrogate data. Dashed lines depict significantly lower relative spindle occurrence compared to surrogate data.

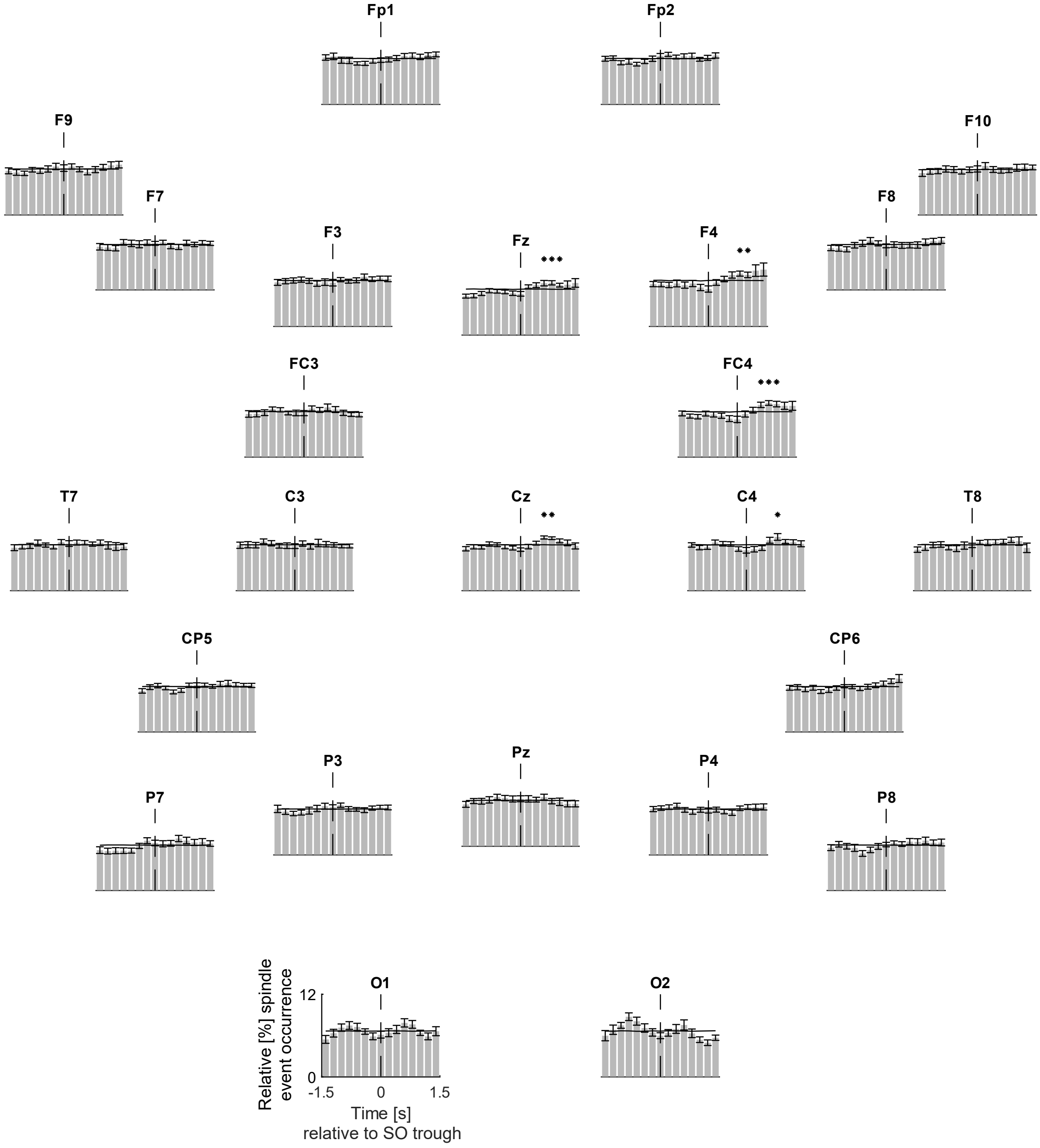


**Figure S3**. Peri-event time histograms of participants younger than one year. Asterisks depict significantly greater relative spindle occurrence compared to surrogate data. Dashed lines depict significantly lower relative spindle occurrence compared to surrogate data.

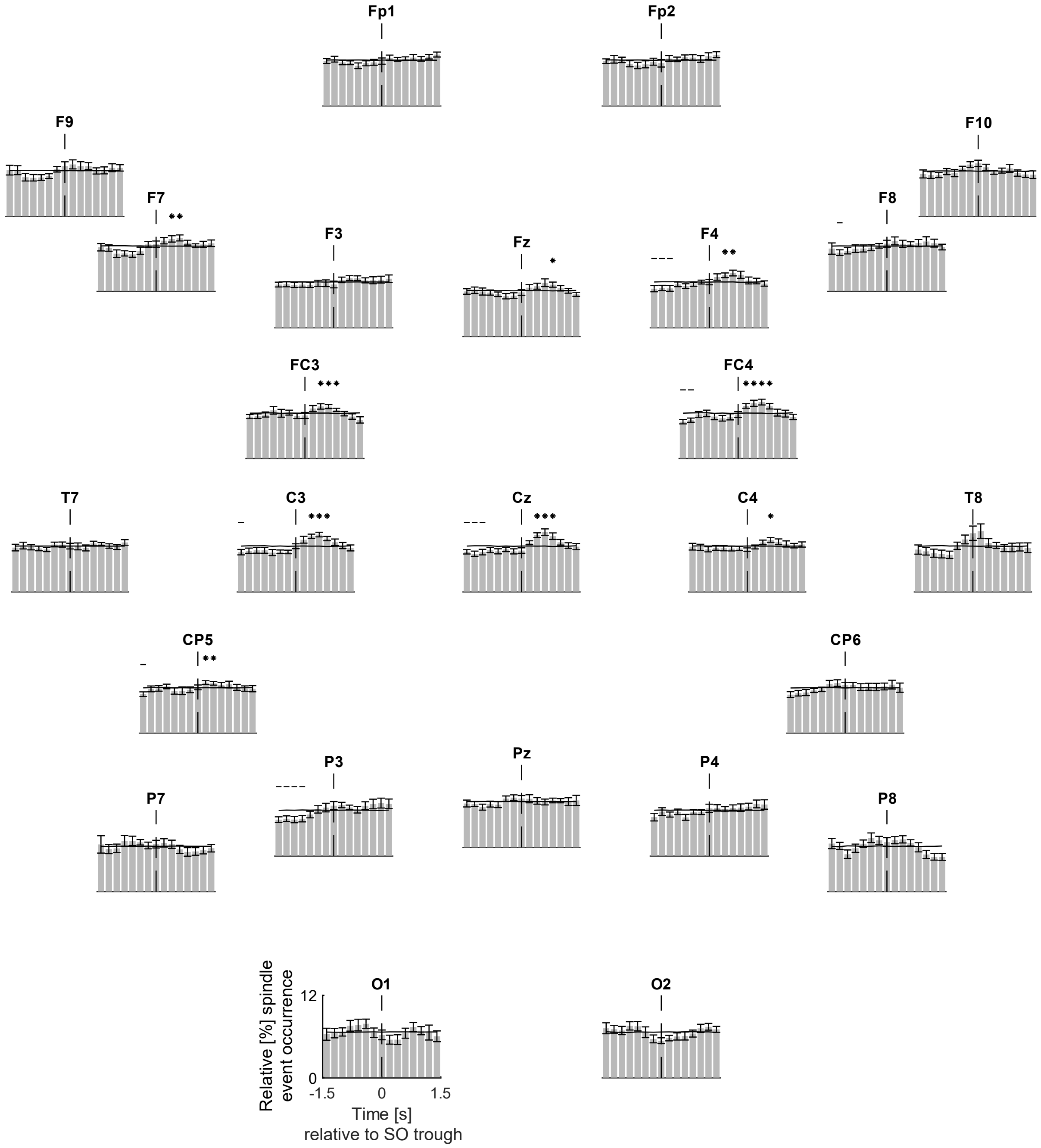


**Figure S3**. Peri-event time histograms of participants older than one year. Asterisks depict significantly greater relative spindle occurrence compared to surrogate data. Dashed lines depict significantly lower relative spindle occurrence compared to surrogate data.

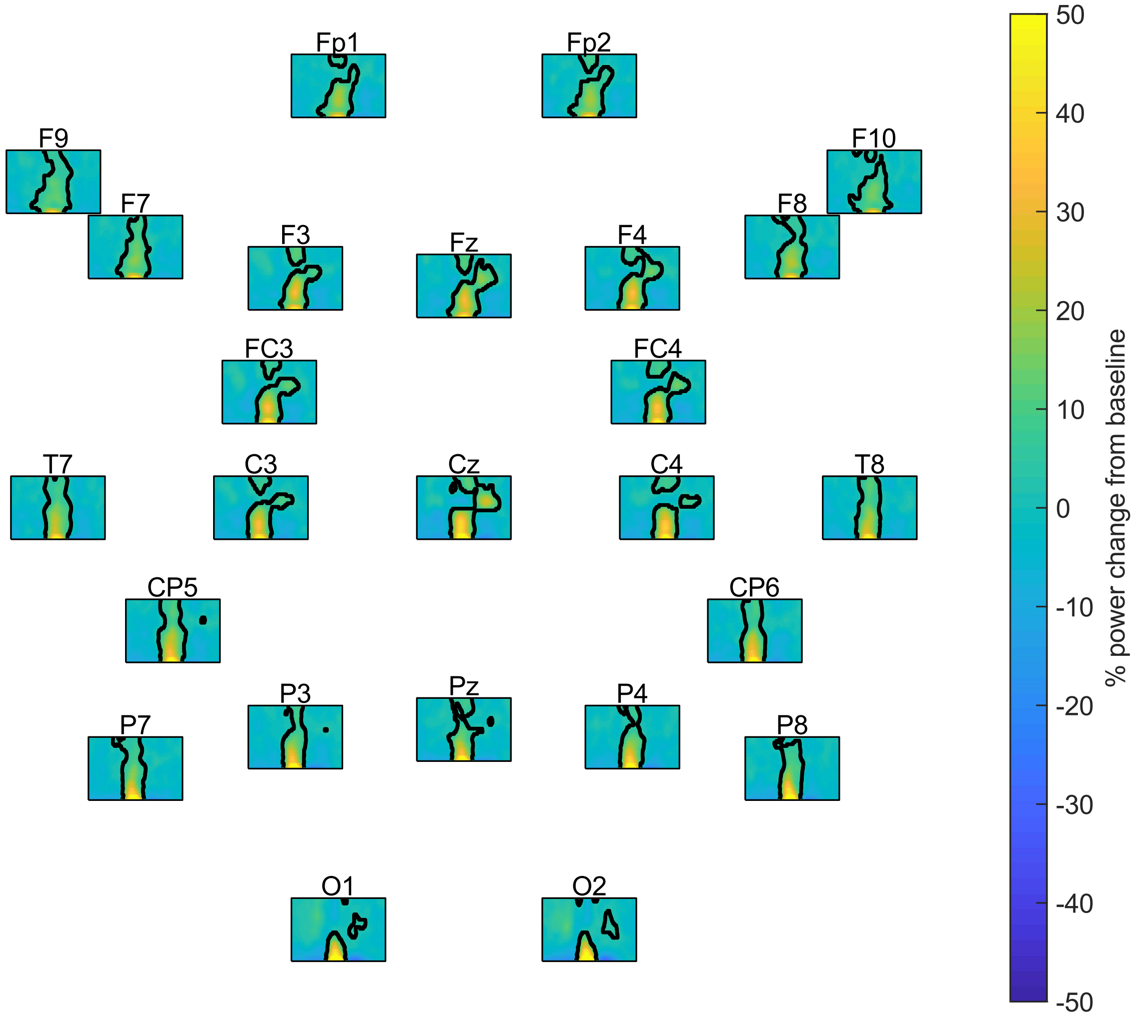


**Figure S4.** Time-frequency representations of SO events across all participants. Outlined areas represent significant changes from baseline. Limits: x-lim: ±1.2 sec around SO trough, y-lim: 3-20 Hz.

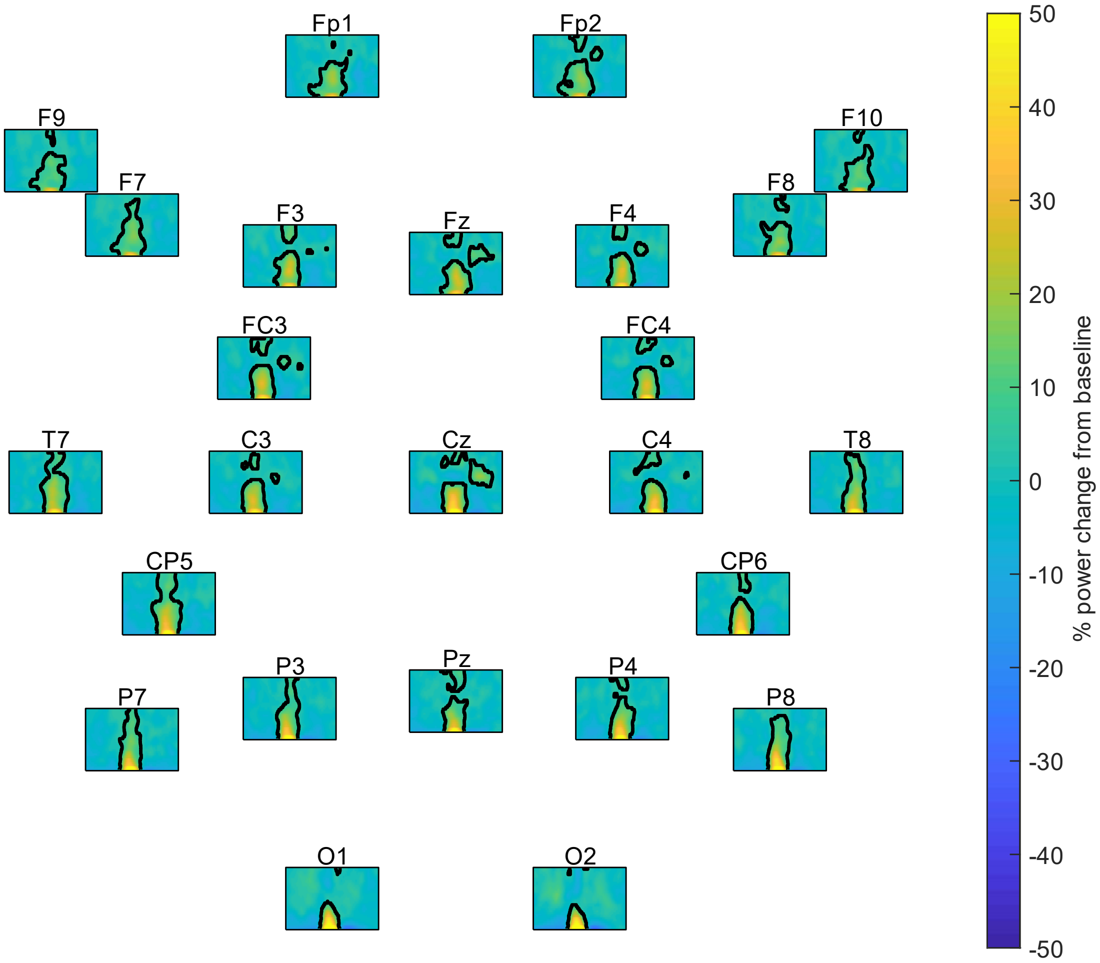


**Figure S5.** Time-frequency representations of SO events of participants younger than 12 months. Outlined areas represent significant changes from baseline. Limits: x-lim: ±1.2 sec around SO trough, y-lim: 3-20 Hz.

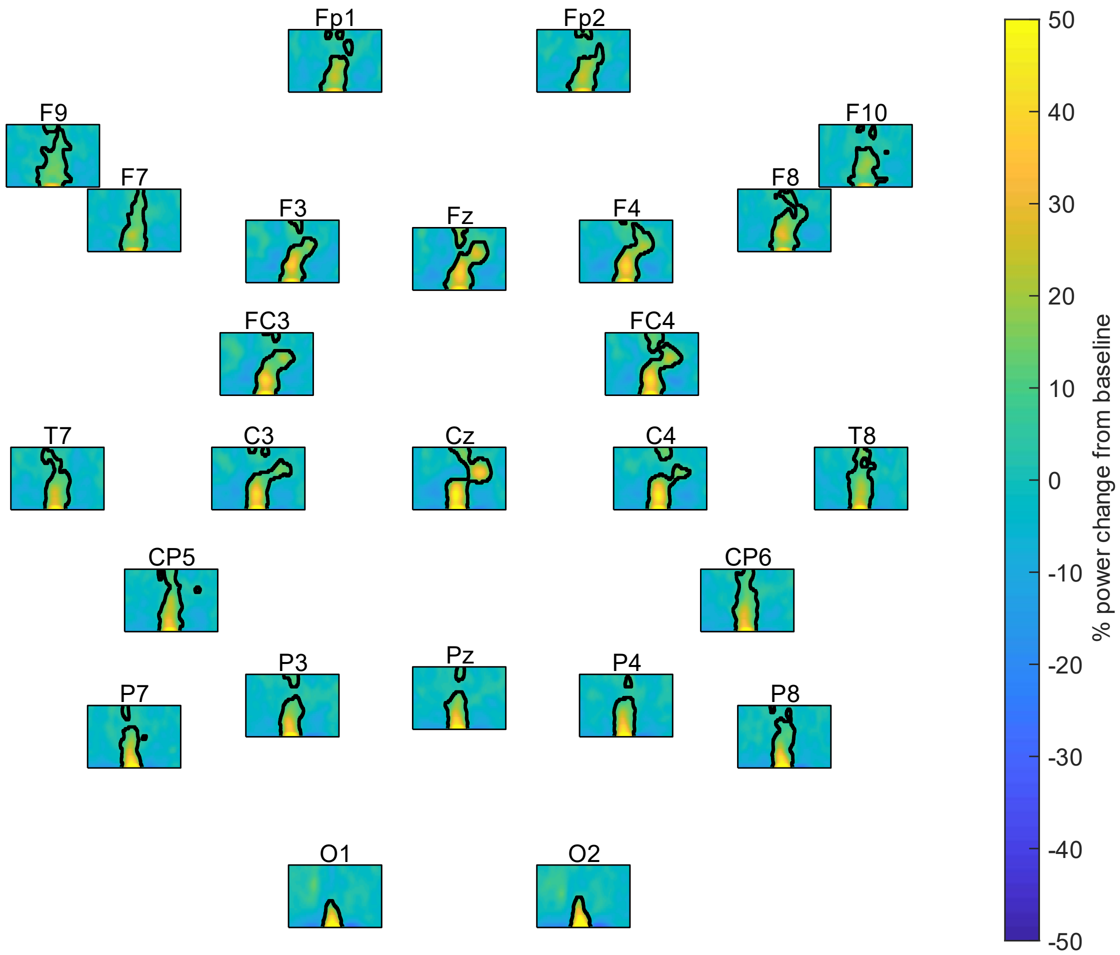


**Figure S6.** Time-frequency representations of SO events of participants older than 12 months. Outlined areas represent significant changes from baseline. Limits: x-lim: ±1.2 sec around SO trough, y-lim: 3-20 Hz.

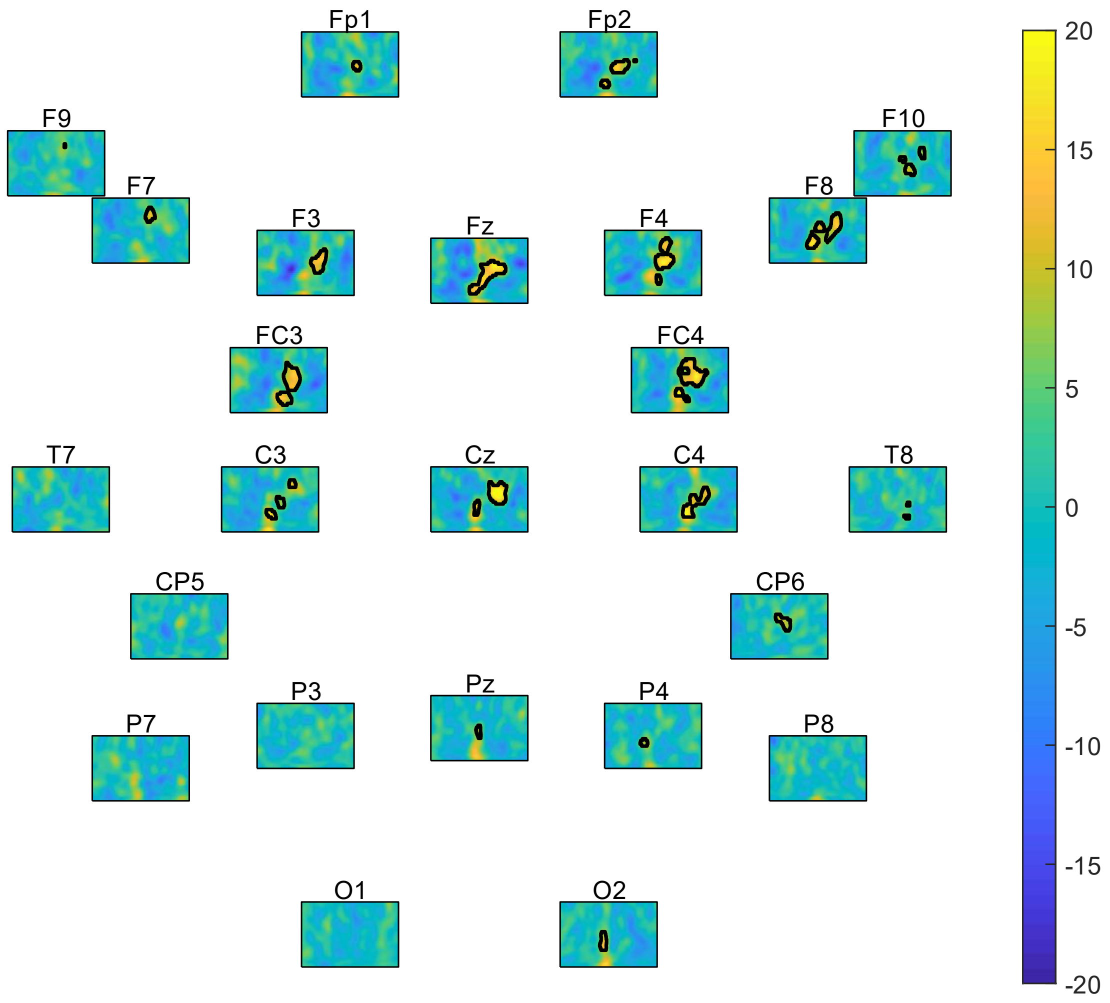


**Figure S7.** Difference in time-frequency representations of SO events of participants older and younger than 12 months. Outlined areas represent significant differences between age groups. Limits: x-lim: ±1.2 sec around SO trough, y-lim: 3-20 Hz.

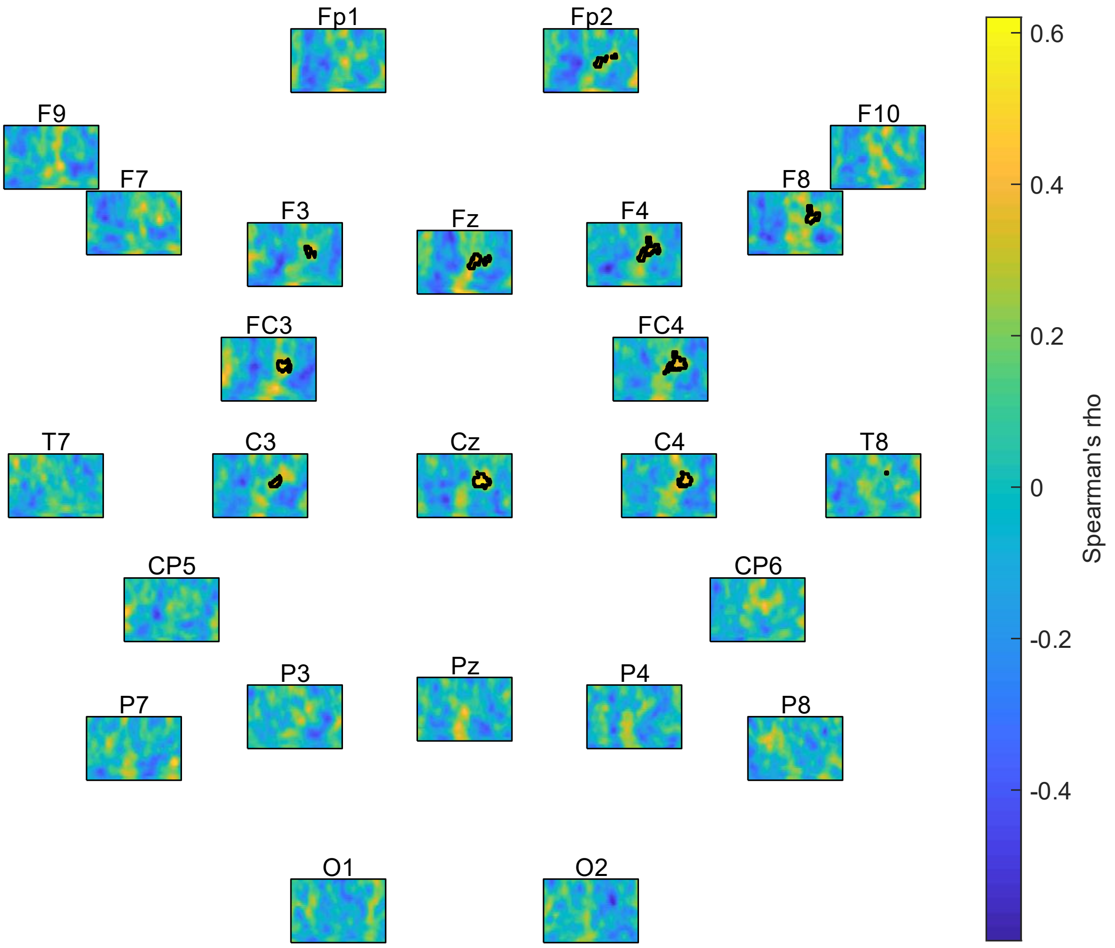


**Figure S8.** Correlation of time-frequency representations of SO events with age (in days). Outlined areas represent significant associations. Limits: x-lim: ±1.2 sec around SO trough, y-lim: 3-20 Hz.

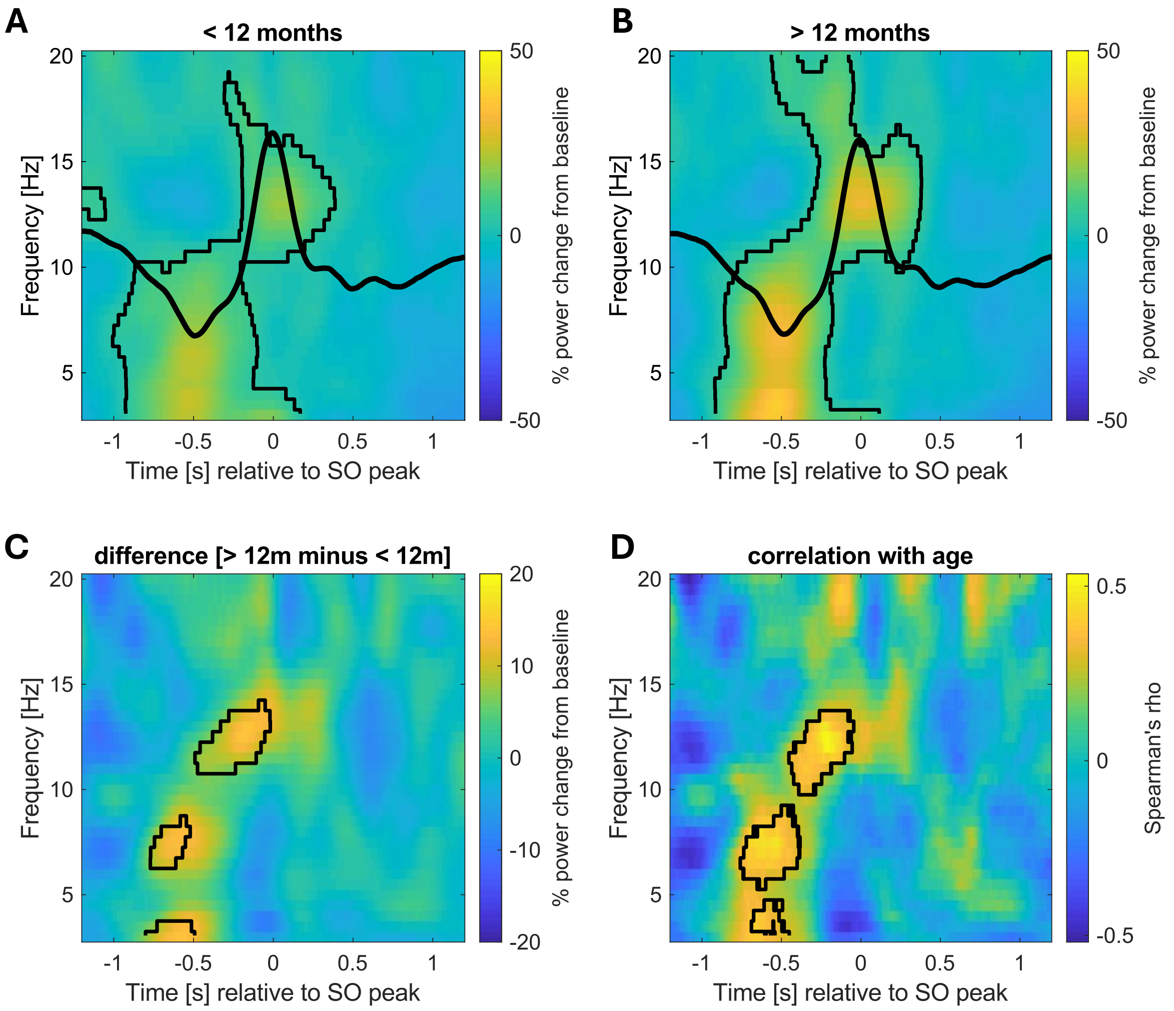


**Figure S9.** Time-frequency representation of SO events locked to the SO peak at exemplary channel Cz in participants younger than 12 months (**A**) and older than 12 months (**B**), as well as the difference between the age groups (**C**). Plots depict the change from baseline (in %). **D**: Correlation of time-frequency representation of SO events with age in days at exemplary channel Cz. Black lines indicate the outline of significant positive clusters. See Figures S10 – S12 for results across all channels.

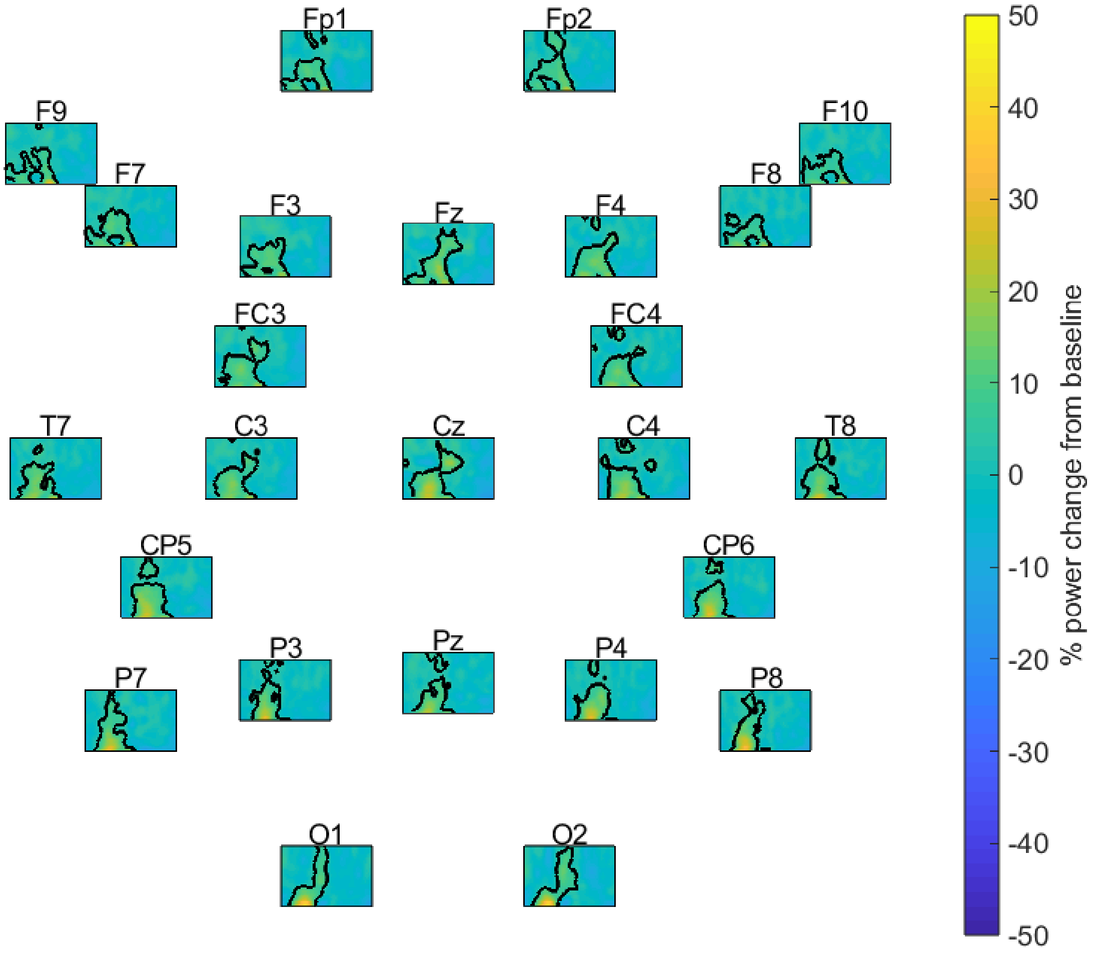


**Figure S10.** Time-frequency representations of SO events (locked to the SO-peak) of participants younger than 12 months. Outlined areas represent significant changes from baseline (cluster $p < .001$). Limits: x-lim: ±1.2 sec around SO peak, y-lim: 3-20 Hz.

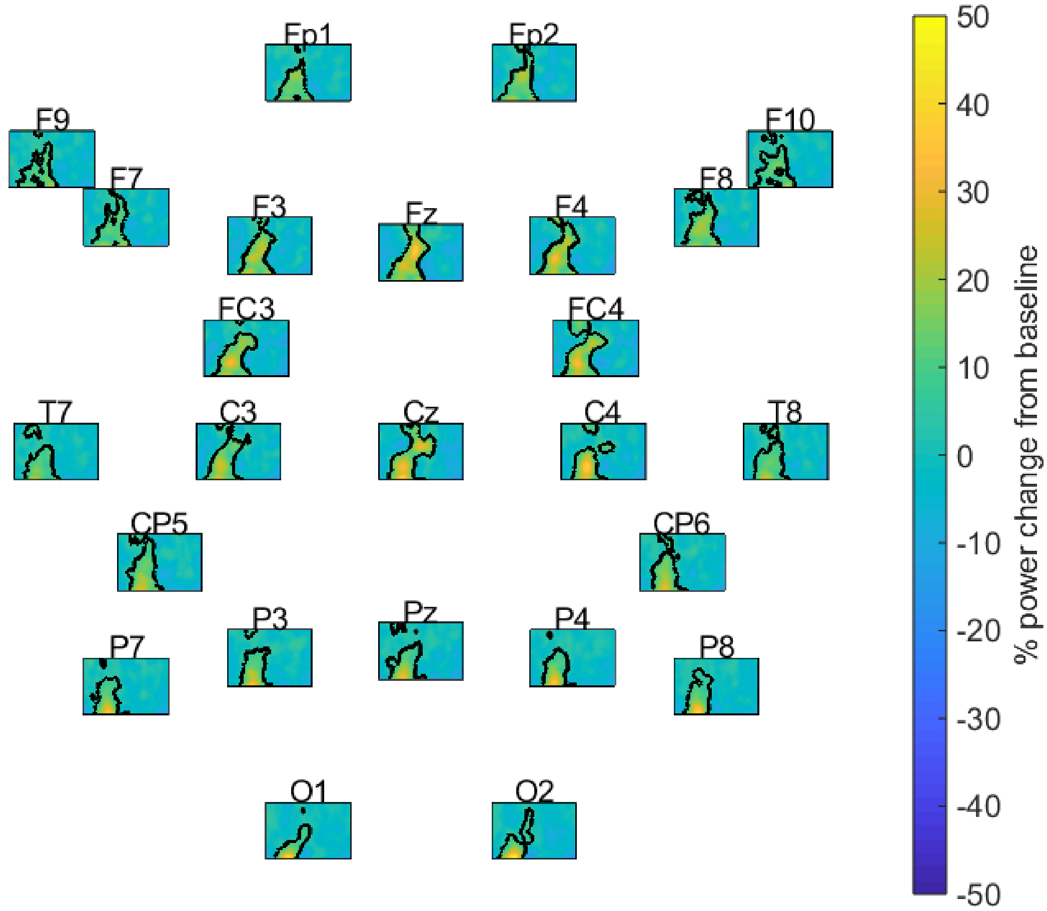


**Figure S11.** Time-frequency representations of SO events (locked to the SO-peak) of participants older than 12 months. Outlined areas represent significant changes from baseline (cluster $p < .001$). Limits: x-lim: ±1.2 sec around SO peak, y-lim: 3-20 Hz.

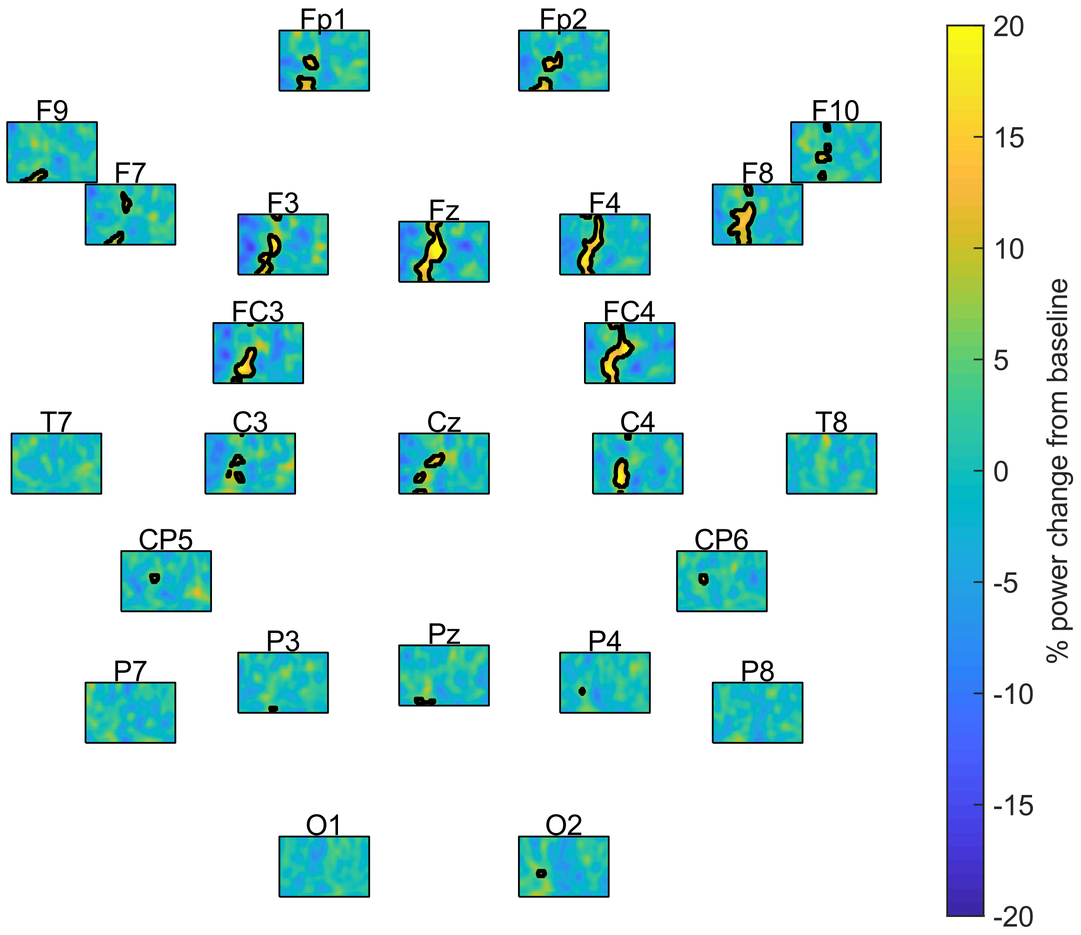


**Figure S12.** Difference in time-frequency representations of SO events (locked to the SO-peak) of participants older and younger than 12 months. Outlined areas represent significant differences between age groups. Limits: x-lim: ±1.2 sec around SO peak, y-lim: 3-20 Hz.
